# Neurogenesis Leads Early Development in Zebrafish

**DOI:** 10.1101/2025.11.12.687769

**Authors:** Zhengduo Wang, Li Tian, Bo Li

## Abstract

Vertebrate early neurogenesis is a highly conserved process fundamental to brain function and the emergence of intelligence. However, the cellular dynamics bridging gastrulation and organogenesis remain elusive due to observational challenges. We developed a live-cell imaging platform for transgenic zebrafish that provides, for the first time, a continuous reconstruction of early neurogenesis across subcellular to organismal scales. Our analysis reveals that neurogenesis is a precisely orchestrated process. Neuronal cell bodies initially coalesce into discrete, linearly arranged clusters extending from the brain along the spinal cord. From these hubs, axons radiate outward to innervate the central nervous system and peripheral tissues, including the yolk sac surface. A primary pioneer neuron projects from the brain, coursing ventrally in parallel to the body axis. Secondary neurons then interconnect, forming a pervasive network that is subsequently refined through selective axonal apoptosis. The emergence of frequent Ca²□ flashes only after structural maturation indicates that functionality is contingent upon an established scaffold. We also observe concurrent material transport and a slow, directional flow of Ca²□ along axons, suggesting complementary signaling modalities. Furthermore, neurogenesis exhibits precise spatiotemporal coupling with histogenesis, particularly with the developing lateral line and vasculature. Our work, with refined spatial and time resolution, defines the kinetic pathway of early neurogenesis and underscores the critical interplay of subsystems in embryogenesis, offering fundamental insights for neural health and bio-inspired intelligence.

## Introduction

Neurogenesis establishes the structural foundation for brain function and organismal behavior (*1–3*). Investigating this process in vivo is therefore essential for understanding the brain and intelligence, and informing artificial system design (*4*). As a critical initiator of embryonic development, neurogenesis is also tightly coordinated with organogenesis and histogenesis (*5*), offering a valuable intervention window for regenerative medicine (*6*). However, the biological events between germ layer formation and organogenesis—the primary neurogenesis period—remain poorly defined, largely due to insufficient dynamic recording at fine temporal resolution. This absence of direct observation spanning cellular and organismal scales impedes a complete understanding of neurogenesis and its developmental interplay.

To elucidate neurogenesis dynamics, we employed long-term in vivo imaging to capture early development in transgenic zebrafish from the zygote stage (**Fig. 1A**). Transgenic lines (**Table S1**) were engineered with fluorescent reporters for neural cells, Ca²□ activity, lateral line, vascular endothelium, slow muscle, and cardiomyocytes (*7–12*), enabling simultaneous visualization of neural architecture, brain function, and other tissues (**Fig. 1B, Fig. S1**). Zygotes were embedded in hydrated agarose under controlled conditions, supporting normal development while preserving all niche conditions. The experimental platform is outlined in **Fig. S2**. The entire 3D processes of neurogenesis and organogenesis were recorded via spinning-disk and two-photon confocal microscopy (**Fig. S3A, Movie S1**). With temporal resolution from 0.5 seconds to 30 minutes and spatial coverage from subcellular to whole-organism scales, we systematically reconstructed neurogenesis, defining its relationship with organogenesis and histogenesis (**Fig. 1C**). Embryos at various hpf underwent transcriptomic analysis, correlating in vivo observations with global gene expression patterns (**Fig. S3B, S3C**).

**Fig. 1.**
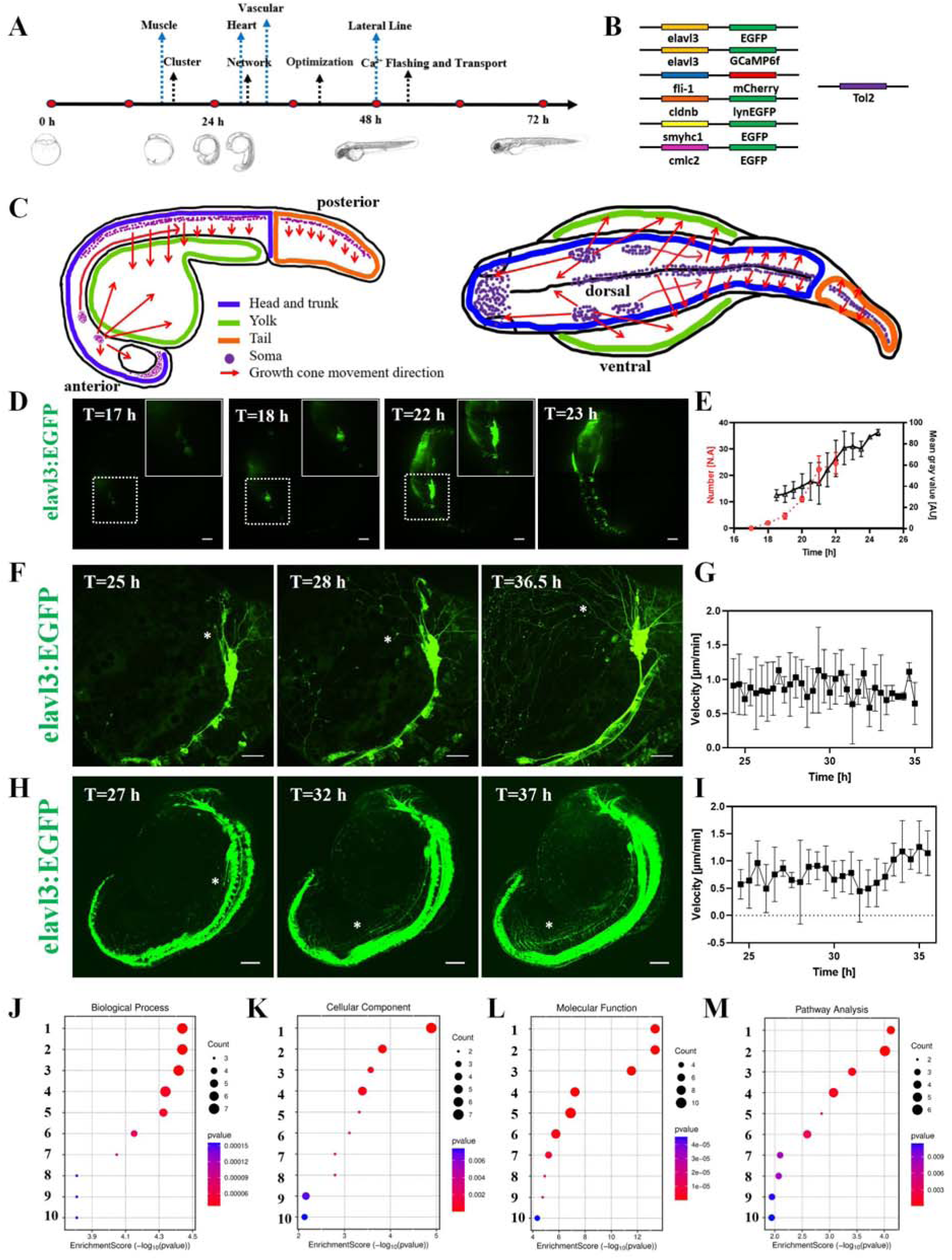
Initial stages of zebrafish neurogenesis. (**A**) Summary of zebrafish embryonic developmental timeline and associated events in this study. The abbreviations for various events are as follows: Cluster: The developmental stage at which neuronal somata begin to aggregate, forming the initial basal structure during early neural development in zebrafish embryos; Network: The developmental stage when the neural network begins to form in zebrafish embryos; Optimization: The developmental stage during neural network formation in zebrafish embryos, characterized by the apoptosis of certain network structures to refine and optimize the overall neural circuitry; Ca2+ Flashing and Transport: The developmental stage in zebrafish embryos when the neural network is initially established, marked by prominent Ca2+ transients in brain neurons and active material transport within the neural network; Muscle: The developmental stage at which distinct skeletal slow muscles become observable in zebrafish embryos; Heart: The developmental stage when cardiac muscle becomes detectable in zebrafish embryos; Vascular: The developmental stage at which blood circulation is observed in zebrafish embryos; Lateral line: The developmental stage when the lateral line neuromasts become visible during zebrafish embryonic development. (**B**) Vectors used for generating transgenic zebrafish lines. All zebrafish employed in this study were generated using the Tol2 transposon system. (**C**) Schematic diagram of early neural development in zebrafish embryos. The left panel shows the lateral view, and the right panel shows the dorsal view. Neuronal somata are depicted in purple, and red arrows indicate the direction of neural network growth. (**D**) Representative confocal image of early neural development in a zebrafish embryo (elavl3:EGFP), with the inset showing a magnified view of clustered neuronal cell bodies. scale bar=200μm. (**E**) The number of clusters (red dashed line) and the optical signal intensity of the predominant brain cluster (black solid line) as a function of time during the initiation of neural development in zebrafish embryos. (**F**) Representative confocal micrographs of early neural network formation in zebrafish (elavl3: EGFP) embryos at 24 hours post-fertilization. Green fluorescence indicates pan-neuronal labeling. The image was acquired using a 30x objective, scale bar=50μm. (**G**) Growth cone velocity as a function of time during neural network development in zebrafish embryos. (**H**) Representative confocal image illustrating neurite outgrowth from the brain toward the spinal cord during neural development in a zebrafish embryo (elavl3:EGFP). scale bar=100μm. (**I**) The growth rate of the primary ventral nerve from the brain to the spinal cord over time during neural development in zebrafish embryos. (**J-M**) KEGG enrichment analysis of neural-related transcriptome data from zebrafish embryos at 12 hpf. (**J**) Biological process terms: 1-10 correspond to chemical synaptic transmission, anterograde trans-synaptic signaling, trans-synaptic signaling, synaptic signaling, regulation of membrane potential, circadian rhythm, regulation of circadian rhythm, glycogen metabolic process, glucan metabolic process, and cellular glucan metabolic process. (**K**) Cellular component terms: 1-10 correspond to postsynapse, postsynaptic membrane, chloride channel complex, synaptic membrane, SMN-Sm protein complex, Cajal body, GABA-A receptor complex, GABA receptor complex, somatodendritic compartment, and spliceosomal complex. (**L**) Molecular function terms: 1-10 correspond to protein threonine phosphatase activity, protein serine phosphatase activity, protein serine/threonine phosphatase activity, phosphoprotein phosphatase activity, phosphoric ester hydrolase activity, phosphatase activity, extracellular ligand-gated ion channel activity, amino acid: sodium symporter activity, amino acid: cation symporter activity, and transmitter-gated ion channel activity. (**M**) Pathway analysis terms: 1-10 correspond to mRNA surveillance pathway, neuroactive ligand-receptor interaction, oocyte meiosis, regulation of actin cytoskeleton, other glycan degradation, adrenergic signaling in cardiomyocytes, vascular smooth muscle contraction, insulin signaling pathway, cellular senescence, and herpes simplex virus 1 infection. For all statistical plots, unpaired Student’s t-test (n > 3 experiments, data are mean ± SD) was conducted for all data. The error bars are the standard error of the mean of the data.

By targeting the developmental period for neural system establishment, which is also the period between germ layer formation and organogenesis, we identified key neurogenesis events and their functions. The initial neural structure comprises two neuronal somata clusters in the brain (**Fig. 1D**, **1E**, **S4A**, **Movie S2**). Axons project from these clusters, forming nascent networks in the brain, yolk sac, and spinal cord (**Fig. 1F**, **1G, S4B**, **Movie S3**). Subsequently, additional clusters emerge caudally, representing spinal cord precursors (**Fig. 1E**). A major ventral neuron (MVN) extends from the brain, coursing parallel to the dorsal spinal cord to innervate somatic regions (**Fig. 1H**, **1I, S4C**, **Movie S4-left**). Hierarchical network assembly, guided by growth cones, dominates neurogenesis (**Fig. 2**, **S5**, **Movie S5**, **S6**). Subsequent structural optimization occurs through selective neuronal apoptosis (**Fig. 3A, 3B, 3C, 3D, 3E, 3F, 3G, 3H**, **S6A**, **S6B**, **S6C, Movie S7**). Finally, network maturation enables functionality, evidenced by delayed onset of Ca²□ transients (**Fig. 3J, 3K**, **Movie S8**) and slow-scale material transport (**Fig. 3M, 3N, 3O, 3P, 3Q, 3R**, **S6D**, **S6E**, **Movie S9**). Furthermore, the MVN exhibits spatial co-localization with the stem cell and lateral line, underscoring the coupling of neurogenesis and histogenesis (**Fig. 4B, 4C, S8B, Table S2**). Blood circulation and organ initiation occur after the neural scaffold forms, highlighting neurogenesis primacy in early development (**Fig. 4H, 4I, 4J, S9**, **Movie S10**, **S11**). Our findings are schematically summarized in **Fig. S11**.

**Fig. 2.**
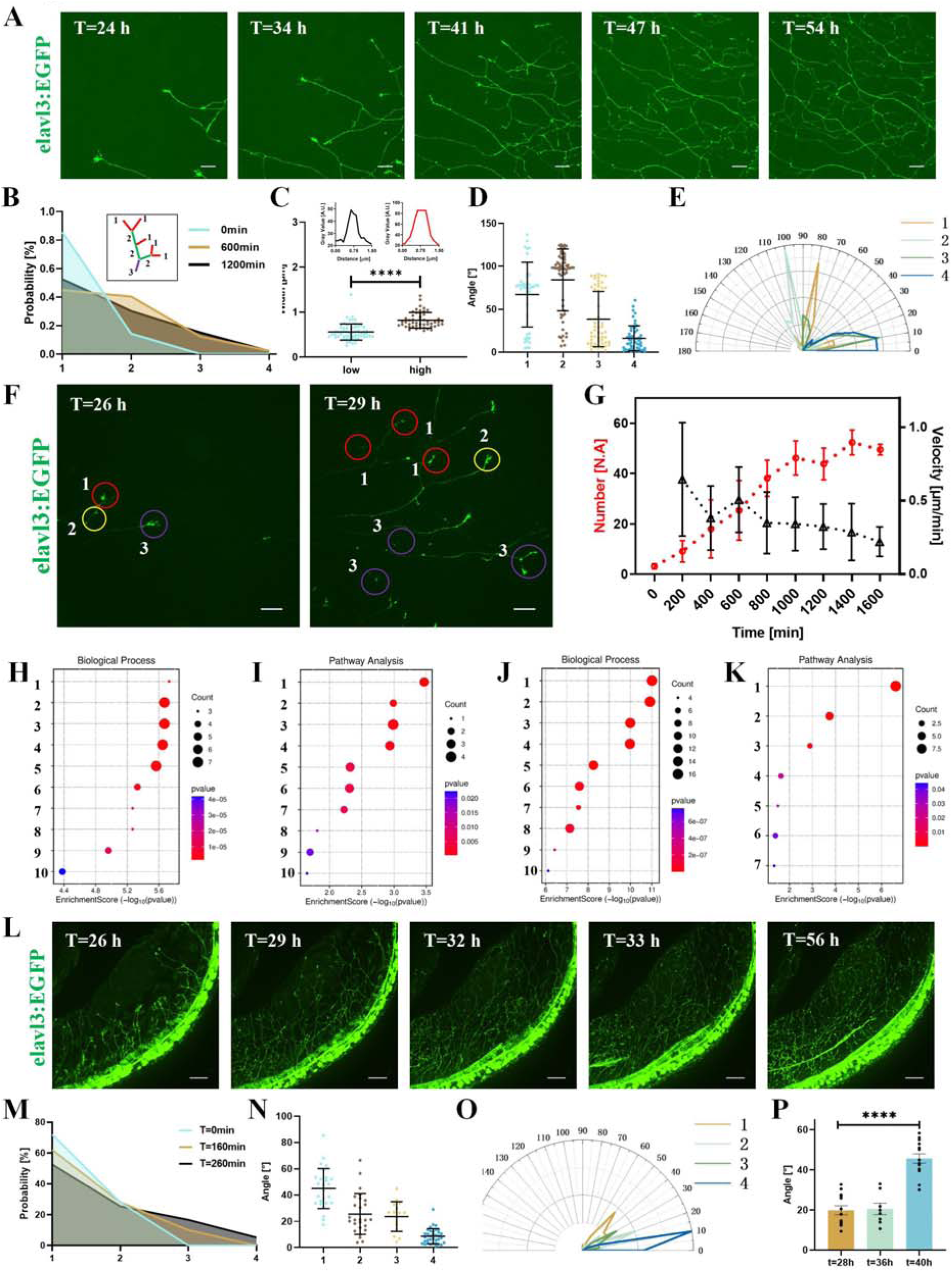
Network formation in the brain and spinal cord regions. (**A**) Representative confocal micrographs of early neural network formation in zebrafish (elavl3: EGFP) embryos at 24 hours post-fertilization. Green fluorescence indicates pan-neuronal labeling. The image was acquired using a 60× objective, scale bar=20μm. (**B**) Frequency distribution of different network hierarchy levels during neural circuit assembly in zebrafish embryos. An axon directly connected to a growth cone is defined as Level 1; Level 2 denotes axons formed by the convergence of two Level 1 axons; Level 3 arises from the merging of two Level 2 axons. When a higher-level and a lower-level axon converge, the resulting axon retains the level of the higher one. (**C**) Quantification of axonal widths across different orders. The inset displays the brightness profiles of low- and high-order axons. (**D**) Statistical analysis of the angle between the direction of movement of growth cones and axons of different hierarchical levels. (**E**) Polar plot showing the probability distribution of the angles between growth cone and axon movement directions. (**F**) Representative confocal image of a growing growth cone, scale bar=20μm. (**G**) Temporal changes in growth cone migration speed (black line) and the number of neural network nodes (red line). (**H, I**) Enrichment analysis of neural-related highly expressed genes in 24 h zebrafish embryos: **H**, Biological process terms: 1-10 correspond to peripheral nervous system neuron axonogenesis, chemical synaptic transmission, anterograde trans-synaptic signaling, trans-synaptic signaling, synaptic signaling, peripheral nervous system development, peripheral nervous system neuron development, peripheral nervous system neuron differentiation, spinal cord development, and central nervous system neuron differentiation, respectively. **I**, Signaling pathway terms: 1-10 represent mRNA surveillance pathway, autophagy-other, neuroactive ligand-receptor interaction, oocyte meiosis, tight junction, adrenergic signaling in cardiomyocytes, Notch signaling pathway, virion-Ebolavirus/Lyssavirus/Morbillivirus, TGF-beta signaling pathway, and glycosphingolipid biosynthesis-ganglio series. (**J, K**) Enrichment analysis of neural-related highly expressed genes in 48 h zebrafish embryos: **J**, Biological process terms: 1-10 correspond to cell projection morphogenesis, cell part morphogenesis, neuron projection morphogenesis, plasma membrane bounded cell projection morphogenesis, axon development, axonogenesis, heterophilic cell-cell adhesion via plasma membrane cell adhesion molecules, cell morphogenesis involved in neuron differentiation, regulation of vascular endothelial growth factor receptor signaling pathway and vascular endothelial growth factor signaling pathway. **K**, Signaling pathway terms: 1-7 represent Neuroactive ligand signaling, Cell adhesion molecules, Virion-Lassa virus and SFTS virus, Notch signaling pathway, Virion-Ebolavirus, Lyssavirus and Morbillivirus, ErbB signaling pathway, and Glycosphingolipid biosynthesis -ganglio series. (**L**) Representative confocal micrograph showing the formation of early neural networks in the spinal cord region of a transgenic zebrafish embryo (Tg(elavl3:EGFP)) at 24 hours post-fertilization (hpf). The green fluorescence indicates pan-neuronal labeling. The top image was acquired with a 30× objective. Scale bar = 50 μm. (**M**) Hierarchy of network integration in the developing spinal cord. The plot shows the distribution of axon ranks, defined by their origin: Rank 1 (direct from growth cones), Rank 2 (from two Rank 1), and Rank 3 (from two Rank 2). Fusion events always preserve the higher rank. (**N**) Angular relationship between growth cone and axon trajectories. The graph statistically compares the direction angles across different hierarchical axon ranks. (**O**) Directional alignment probability. The polar plot illustrates the distribution of angles between the movement vectors of growth cones and axons. (**P**) Statistical analysis of the angles between the network extending from neuronal somata adjacent to the spinal cord and the direction of growth cone movement at different time points during spinal neural network development. For all statistical plots, unpaired Student’s t-test (n > 3 experiments, data are mean ± SD) was conducted for all data with **** P < 0.0001, *** P < 0.001, ** P < 0.1, * P <0.5, ns P > 0.5. The error bars are the standard error of the mean of the data.

**Fig. 3.**
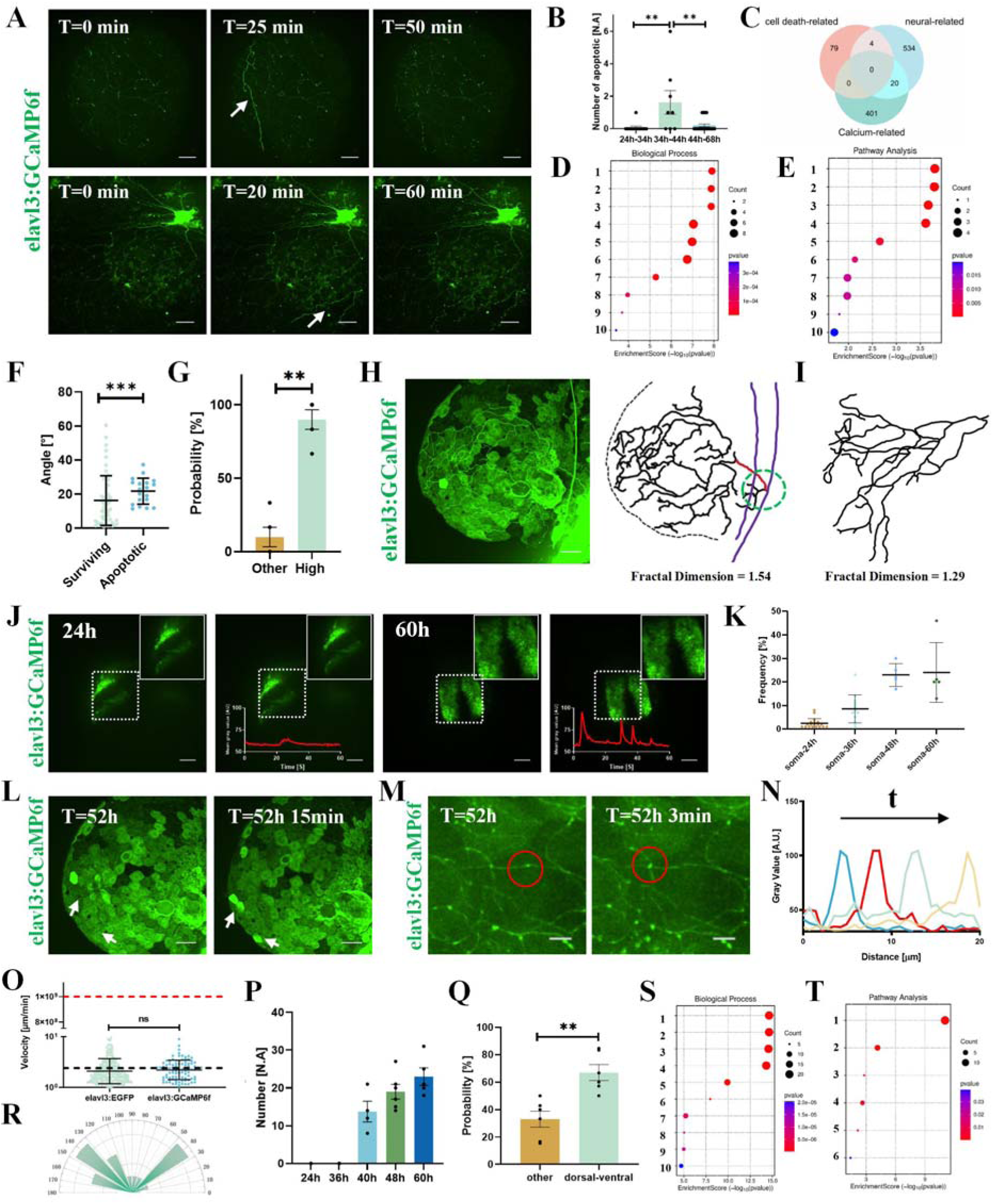
Apoptosis assisted network optimization and brain function after network maturation. (**A**) Representative confocal images showing the refinement of a subset of axonal networks via apoptosis during neural circuit formation in zebrafish (elavl3: GCaMP6f) embryos. White arrows indicate pruned axons. Scale bar = 50μm. (**B**) Quantification of apoptosis events at different developmental time points in zebrafish embryos. (**C**) Venn diagram showing the overlap among highly expressed neuro-related, calcium-related, and cell death-related genes from transcriptome data analysis of zebrafish embryos. (**D, E**) Enrichment analysis of highly expressed death-related genes in 60 hpf zebrafish embryos. **D**, Biological process terms:1-10 correspond to positive regulation of apoptotic process, positive regulation of programmed cell death, positive regulation of cell death, regulation of apoptotic process, regulation of programmed cell death, regulation of cell death, apoptotic signaling pathway, extrinsic apoptotic signaling pathway,MyD88-dependent toll-like receptor signaling pathway, and extrinsic apoptotic signaling pathway via death domain receptors. **E**, Signaling pathway terms:1-10 represent Herpes simplex virus 1 infection, apoptosis, cytokine–cytokine receptor interaction, autophagy–animal, necroptosis, p53signaling pathway, Salmonella infection, cytoskeleton in muscle cells, virion – Ebolavirus/Lyssavirus/Morbillivirus, and MAPK signaling pathway. (**F**) Statistical analysis of the angles between apoptotic versus non-apoptotic neurons and the axonal network in the zebrafish neural network. (**G**) Probability of neuronal apoptosis events involving neurons of hierarchy level >2 in the developing zebrafish embryonic neural network. (**H**) Representative confocal image (left) and corresponding structural diagram (right) of large-scale neural network apoptosis in a relatively mature zebrafish embryonic neural network. Red and green dashed circles indicate locations of the highest hierarchy axons; purple lines denote higher-level axonal projections. (**I**) Representative structure of a non-apoptotic neural network. (**J**) Representative confocal images showing calcium transients in a zebrafish (elavl3:GCaMP6f) embryo at 24 hpf (left) and 60 hpf (right), the right panel shows magnified views. Line chart showing the dynamics of cerebral neuronal calcium signals in zebrafish embryos at 24 and 60 hpf. Scale bar = 50μm. (**K**) Statistical analysis of calcium flash frequency at different developmental stages of zebrafish embryos. (**L**) Representative confocal image showing calcium transients observed on the yolk sac surface of a zebrafish embryo (elavl3:GCaMP6f) at 60 hpf. Scale bar=50μm. (**M**) Representative confocal images showing putative material transport within the axonal network of zebrafish (elavl3: GCaMP6f) embryos at approximately 48 hpf. Arrows indicate observed transported cargo. (**N**) This figure shows the positions of material transport within the same neural axon network at different time points. The positions of the substances are represented by grayscale values. (**O**) Quantification of cargo transport velocity within the axonal network of zebrafish embryos (elavl3: GCaMP6f; elavl3: EGFP). The black dashed line denotes the average velocity of growth cone migration, and the red dashed line represents the average signal conduction velocity in myelinated axons. (**P**) Quantification of cargo transport duration per unit time observed in the neural network at different time points during zebrafish embryonic development. (**Q**) Bar graph showing the probability of observed cargo transport directions within the neural network, comparing dorsal-to-ventral transport against all other directions. (**R**) Polar histogram of cargo transport direction probabilities in brain. (**S, T**) Enrichment analysis of neural-related highly expressed genes in 60 hpf zebrafish embryos. **S**, Biological process terms: 1-10 correspond to chemical synaptic transmission, anterograde trans-synaptic signaling, trans-synaptic signaling, synaptic signaling, regulation of membrane potential, cholinergic synaptic transmission, positive regulation of developmental process, chloride transmembrane transport, chloride transport, and positive regulation of cell differentiation. **T**, Signaling pathway terms: 1-6 represent Neuroactive ligand-receptor interaction, Cell adhesion molecules, Glycosphingolipid biosynthesis – ganglio series, Oocyte meiosis, Autophagy-other, and VEGF signaling pathway. For all statistical plots, unpaired Student’s t-test (n > 3 experiments, data are mean ± SD) was conducted for all data with **** P < 0.0001, *** P < 0.001, ** P < 0.1, * P <0.5, ns P > 0.5. The error bars are the standard error of the mean of the data.

**Fig. 4.**
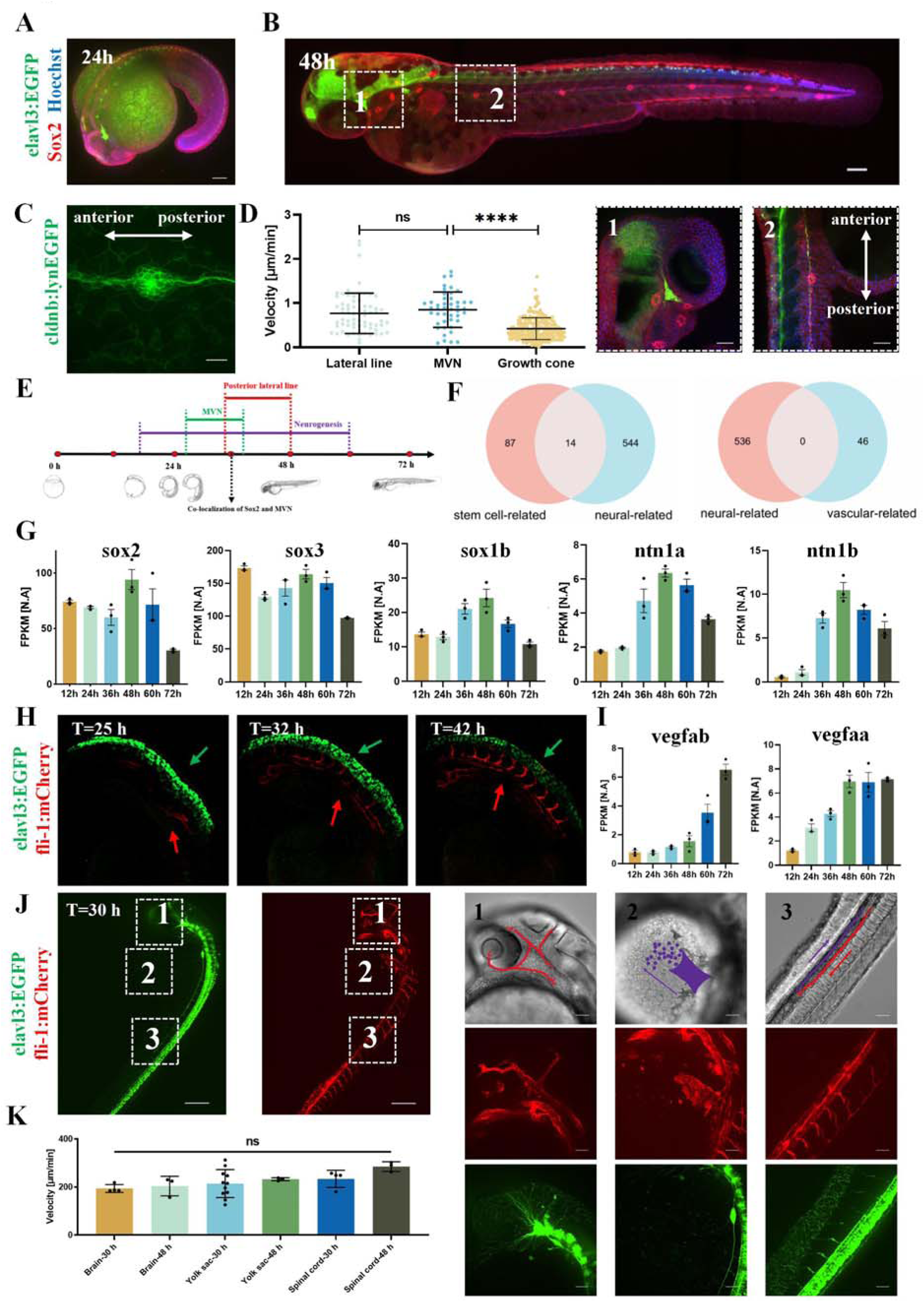
Coupling between neurogenesis with histogenesis and organogenesis. (**A**) Representative immunofluorescence images of the nervous system and Sox2 in zebrafish embryos (elavl3:EGFP) at 24 hours of development. Sox2 is labeled in red, nuclei are counterstained with Hoechst (blue), and pan-neuronal labeling is shown by EGFP signal (green) in the transgenic line. Scale bars =100 µm. (**B**) Immunofluorescence analysis of the nervous system and Sox2 in zebrafish embryos (elavl3:EGFP) at 48 h. Scale bar =100 µm. The inset below shows a high-resolution confocal image of the white-boxed region in Panel B. Scale bars =50 µm. The color scheme is identical to that in A. (**C**) Representative confocal images of the posterior lateral line in zebrafish (cldnb:lynEGFP) embryos at 48 hpf. Scale bar =20 µm. (**D**) Quantification of the migration speed of the posterior lateral line primordium, the migration speed of major ventral neurons (MVN) along the spinal cord, and the growth cone speed during zebrafish embryonic development. (**E**) Timeline of neurogenesis, MVN formation, and posterior lateral line development during zebrafish embryogenesis. (**F**) Venn diagrams showing the overlap between neuro-related genes and stem cell-related genes (top), as well as between neuro-related genes and vascular-related genes (bottom), based on transcriptomic data. (**G**) Normalized expression of Sox2, Sox3, Sox1b, Ntn1a, and Ntn1b genes at different developmental time points of zebrafish embryos from transcriptome data. (**H**) Representative bright-field image of early neural and vascular development in a zebrafish embryo. Green fluorescence labels pan-neuronal structures; red fluorescence marks vascular endothelium. Green arrows indicate neuronal signals; red arrows indicate vascular signal. (**I**) Normalized expression of Vegfab and Vegfaa genes at different developmental time points of zebrafish embryos from transcriptome data. (**J**) Representative confocal images showing nervous and vascular development in a zebrafish embryo at 30 h, obtained from a cross between elavl3:EGFP and fli1a:mCherry transgenic lines. The right panels display bright-field, vascular (mCherry), and neuronal (EGFP) images of the brain, yolk sac, and spinal cord regions. The leftmost confocal image was acquired with a 40x objective lens. (**K**) Statistical Analysis of Blood Cell Flow Velocity in the Brain, Yolk Sac, and Spinal Cord of Zebrafish Embryos at 30 h and 48 h of Development. The three sets of images on the right were obtained using a 30x objective lens. Scale bars =50 µm. For all statistical plots, unpaired Student’s t-test (n > 3 experiments, data are mean ± SD) was conducted for all data. The error bars are the standard error of the mean of the data.

### The initial stage of neurogenesis exhibits highly organized features

Neural system development is spatiotemporally segregated into brain and spinal cord regions. To establish a unified timeline, we define t = 0 as the fertilization time point, maintained consistently throughout this study. Neural development initiates first in the brain region. By twelve hours post-fertilization, several neuronal somata coalesce, constituting the first organized structure of the nervous system (**Fig. 1D**, **S4A**, **Movie S2**). Two cell clusters emerge at the anteriormost aspect of the embryo, positioned bilaterally in the brain. The position of the cell clusters corresponds to the transient area between the midbrain and hindbrain, but is distal to the forebrain in the adult fish. The fluorescence intensity of these clusters increases dramatically around 20 hours (**Fig. 1E**), indicating a rapid expansion in neuronal number.

Axons subsequently extend from these clustered somata, forming an interconnected network that encompasses both the brain and yolk sac regions (**Fig. 1F**, **S4B**, **Movie S3**). Notably, the yolk sac constitutes an integral component of this early neurogenesis. Rather than developing independently, extending axons first elaborate within the brain before projecting to colonize the yolk sac surface, ultimately forming a continuous network (**Fig. 1F**, **S4B**, **Movie S3**). Growth cones maintain a consistent extension rate of 0.9 µm/min throughout this process (**Fig. 1G**). The transgenic zebrafish line utilizes the elavl3 promoter to drive EGFP expression. As elavl3 is a specific pan-neuronal marker, the observed signals unequivocally identify neural structures, excluding non-neural tissues such as epithelium. This confirms the yolk sac is actively innervated by brain-derived axons during neurogenesis.

Spinal cord neurogenesis initiates nearly concurrently with brain development. Shortly after initial brain clusters form, additional clusters emerge along the embryonic axis, establishing the spinal cord framework (**Fig. 1F, S4B**). These dorsally positioned structures are designated dorsal cell body clusters (DCBCs). Rather than appearing sequentially, DCBCs emerge synchronously at t = 20 hours (**Fig. 1E, S4B, Movie S3**), indicating coordinated development that precludes anterior-to-posterior signaling. Subsequently, a growth cone projects ventrally from the brain cluster, extending caudally at constant velocity (**Fig. 1H, 1I, S4C, Movie S4-left**), establishing the major ventral neuron (MVN) parallel to dorsal DCBCs. Signaling thus proceeds along the anterior-posterior axis. Finally, branching axons interconnect DCBCs and the MVN with precise somite correspondence, as signaling converts to the dorsal-ventral axis. These observations establish DCBCs and the MVN as the structural framework for spinal cord neural architecture.

Transcriptomic analysis corroborates the *in vivo* observations (**Fig. 1J, 1K, 1L, 1M)**. Biological process enrichment underscores early synaptic signaling, while cellular component analysis reveals foundational neural structures and GABAergic receptor complexes (**Fig. 1J, 1K**). Molecular function highlights serine/threonine phosphatase, ion channel, and transporter activities (**Fig. 1L**). Pathway analysis confirms the prominence of initial synaptic formation (**Fig. 1M**). Collectively, these KEGG results demonstrate that by 12 hpf, zebrafish neural development reaches the foundational stage, preparing functional activation.

### Network establishment dominates neural development

Neurogenesis is governed by the establishment and maturation of axonal networks extending from neuronal somata clusters. In both brain and spinal cord regions, select growth cones exhibit directed migration. These pioneer growth cones establish major axons that form the primary neural framework (**Fig. 2A**, **2L, S5**, **Movie S5**, **S6**). Secondary axons subsequently branch from these primary pathways, elaborating the network complexity (**Fig. 2A**, **2L, S5**, **Movie S5**, **S6**). In the brain, major axons project from initial clusters to other brain regions or the yolk sac (**Fig. 2A**, **S5A**, **Movie S5**), whereas spinal cord axons extend along the dorsal-to-ventral axis (**Fig. 2L**, **S5B**, **Movie S6**). No discernible movement of neuron soma was observed during the network establishment, being consistent with the behavior of glia cells (*13*) and distinguishing the neurogenesis from the migration of stem cells (*14*). In contrast to angiogenesis, in which blood vessels appear simultaneously (**Fig. 4H**, **Movie S10**), the establishment of a neural network involves free exploration of growth cones and extension of axons (**Fig. 2A**, **Movie S5**, **S6**). This fundamental difference in the kinetic pathway suggests that the network structure resulting from neurogenesis offers guidance to angiogenesis and leads to subsequent organogenesis.

To correlate growth cone dynamics with network topology, we implemented Strahler number analysis (*15*) for brain and yolk sac axons. Higher Strahler values indicate greater axonal convergence (major pathways), while lower values represent terminal branches. The evolving probability distribution of Strahler values quantitatively captures network maturation (**Fig. 2B**). Anatomically, higher-order axons exhibit larger diameters (**Fig. 2C**), consistent with observations in the brain of Drosophila Melanogaster (*16*). Structurally, major axons maintain smaller angular deviations from growth cone trajectories than branching axons (**Fig. 2D**), reflecting functional specialization. Intermediate Strahler values, namely rank 2 and 3, display bimodal angular distributions (**Fig. 2E**), suggesting their transitional role in network connectivity. Dynamically, growth cones exhibit directed migration rather than stochastic exploration (**Fig. 2F**), resulting in highly aligned major axon orientation. Concurrently, growth cone velocity decays significantly over time (**Fig. 2G**), consistent with chemotactic mechanisms dependent on chemical gradients. Neural network establishment coincides with node number saturation and negligible growth cone speeds (**Fig. 2G**).

By 24 hpf, the transcriptomic profiles of neural development shifted to specialized maturation, with analyses highlighting neuronal differentiation in central and peripheral nervous systems and strengthened synaptic signaling (**Fig. 2H, 2I**). By 48 hpf, development progressed to structural refinement and functional maturation, marked by precise axonal morphogenesis and synaptic transmission, with pathway enrichment revealing upregulation of axon development and guidance processes (**Fig. 2J, 2K**).

The kinetic pathway of network establishment in the spinal cord closely mirrors that of the brain. In concert with the caudal extension of the major ventral axon, multiple growth cones emerge from dorsal cell clusters, exhibiting dorsal-to-ventral trajectories (**Fig. 2L**, **S5B**, **Movie S6**). Secondary axons with more stochastic orientations subsequently elaborate the network (**Fig. 2M**, **2N**, **2O**). Notably, as the MVN elongates, other major axons reorient from strict dorsal-to-ventral alignment to trajectories parallel with the MVN (**Fig. 2P**). This reorientation suggests these axons establish connectivity with the MVN and are guided by its extension - a kinetic pathway absent in the brain-yolk sac system due to the lack of an analogous migratory traction force or signaling source.

### Structural pruning by apoptosis precedes neural function

During late network establishment, *in vivo* observation and transcriptomic data reveal frequent apoptosis in specific brain and yolk sac neurons. Upon apoptosis, intense Ca²□ transients propagate throughout the axon (**Fig. 3A, S6A-C, Movie S7-left**). Apoptosis frequency peaks at 36 hpf (**Fig. 3B**), coinciding with network consolidation and suggesting a role in optimization. Transcriptomic data show concurrent upregulation of genes related to cell death, neurogenesis, and Ca²□ signaling, though no gene spans all three categories (F**ig. 3C****, Table S3, S4**). Ca²□ signaling has been implicated in apoptosis in cell lines (*17, 18*); our *in vivo* observations confirm this and suggest its function is axonal network pruning. Pathway enrichment confirms structural refinement and functional adaptation, with enhanced axonal morphology and synaptic precision (**Fig. 3D, 3E**). We hypothesize that apoptosis refines the network by eliminating suboptimal connections, maximizing information transfer efficiency.

To test this, we analyzed the characteristics of neurons undergoing apoptosis. Pruned neurons consistently exhibit two features: a higher Strahler order and a significant deviation in their orientation from the original growth cone trajectory (**Fig. 3F, 3G**). Following network completion, apoptosis persists at a lower frequency (**Fig. 3B, 3H**, **Movie S7-right**). However, neurons eliminated at this stage are markedly more mature. As exemplified in Fig. 4h, the Ca²□ transient can propagate across the entire yolk sac via a single, intricately branched neuron. Compared with normal neurons with hierarchical features (**Fig. 3I**), those that underwent apoptosis have more complex geometrical features, including a higher value of fractal dimension. The loss of such mature neurons induces more extensive structural compromise and carries a higher metabolic cost, which partly explains the observed reduction in apoptosis rate after network finalization.

Beginning at 36 hours post-fertilization, Ca²□ transients initiate within neuronal somata (**Fig. 3J**, **Movie S8**), with their frequency increasing over time (**Fig. 3K**). As Ca²□ is a key mediator of neural signaling and a hallmark of neuronal activity (*19*), its emergence only after network structural maturity supports the principle that function is contingent upon established architecture. Notably, these Ca²□ transients occur not only in the brain but also across the yolk sac surface, which is now populated with neuronal cell bodies (**Fig. 3L**). Given the absence of elavl3 signal in the yolk sac at 20 hpf (**Fig. 1D**), these neurons cannot be yolk sac-native. Since axons previously projected from brain clusters to innervate the yolk sac (**Fig. 1F**), we thus posit a testable hypothesis that these somata result from neuronal migration from the brain.

Besides somatic Ca²□ transients, we observe mass transport along axons. Particulate structures, slightly larger than axonal diameters, move bidirectionally within axonal tracts (**Fig. 3M**, **3N, S6D, S6E**, **Movie S9**) at velocities orders of magnitude slower than electrical signals (*20*) or calcium sparks (*21*), identifying a novel transport phenomenon. Its occurrence in both elavl3:EGFP and elavl3:GCaMP6f lines at similar speeds (**Fig. 3O**) suggests these moving entities are highly localized Ca²□ waves. The number of transported particles increases dramatically (**Fig. 3P**), mirroring the rise in Ca²□ transient frequency. The direction of transportation is not random: anisotropic in the brain (**Fig. 3R**) and predominantly dorsoventral in the spinal cord (**Fig. 3Q**). This implies that chemotactic gradients guide not only growth cones (*22*) but also material transport. By 72 hpf, the neural system reaches a state of functional maturation and homeostasis, with fully developed structure and activity (**Fig. 3S, 3T**).

### Interplay between neurogenesis with histogenesis, and organogenesis

Neurogenesis proceeds in concert with histogenesis, the process of cellular differentiation into tissues. To delineate their relationship, we simultaneously visualized biomarkers for neurogenesis and stem cell differentiation via immunostaining for Sox2 (stemness), cldnb (cellular mechanics), and elavl3 (neurogenesis) in embryos at various post-fertilization timepoints. Sox2 is known to be highly expressed in stem cells during early embryonic development (*23*). In our zebrafish embryos, Sox2 is strongly expressed at the animal pole at 3 hpf (**Fig. S7A**). After the onset of neural development, the distribution of the Sox2 signal shows complex spatiotemporal coupling with that of the elavl3 signal (**Fig. 4A, 4B, S7**).

At 48 hpf, immunostaining reveals strong spatial co-localization of elevated Sox2 with the MVN (**Fig. 4B**) and lateral line (cldnb) (**Fig. S8B**). A longitudinal Sox2^+^ domain aligns precisely with the MVN tract, interconnecting spherical rosette-shaped cell clusters (**Fig. 4B, S7E, S8B**). These clusters localize to the brain, yolk sac periphery, and spinal cord (**Fig. 4B, S7**). This co-localization persists from 48 hpf (**Fig. 4B, S7E, S8B**), demonstrating sustained synergy between neurogenesis and stem cell differentiation. As Sox2 coordinates stemness maintenance (*23*) and cluster formation (*24*), our observations reveal a spatially organized interplay between Sox2 and neural development. Notably, Sox2 begins at 3 hpf, preceding neural differentiation, suggesting histogenesis may regulate early neurogenesis (**Fig. S7A, S7B**). Conversely, organized Sox2^+^ domains appear only after MVN and spinal cord maturation, indicating neurogenesis subsequently directs histogenesis (**Fig. 4B, S7E, S8B**).

Anatomically, MVN co-localizes with the lateral line, indicating neurogenesis directs mechanosensory establishment (**Fig. 4C, S7E, S8B, Movie S4-right**). Sox2^+^ bulges completely overlap with lateral line clusters responsible for force sensing (*25*). Growth cone migration speeds during lateral line and MVN formation are significantly higher than during network establishment (**Fig. 4D**). Our *in vivo* observations provide direct evidence that Sox2 mediates lateral line development, corroborating earlier reports (*26, 27*). The sequential appearance of elavl3, Sox2, and cldnb at identical positions suggests a pheromone-like model (*28*) in which earlier-expressed markers direct subsequent patterning. In contrast, actin shows markedly weaker spatial correlation with other markers (**Fig. S8C**). As a cytoskeletal protein (*29*), its ectopic pattern suggests regulation by Sox2 rather than a driving role—supported by molecular evidence (*30*).

Developmental timeline analysis revealed that neurogenesis starts at approximately 16 hpf and continues until prominent functional activity of brain neurons (calcium flashes) is observed at 60 hpf. The MVN begins to form at around 27 hpf and completes by approximately 38 hpf, whereas the posterior lateral line initiates at about 34 hpf and persists until 48 hpf (**Fig. 4E**). Their temporal expression sequences validate our analysis at the transcriptomic level — genes predominantly responsible for stem cell differentiation precede those primarily governing neuronal growth (**Fig. 4F, 4G, Table S5**).

Subsequently, blood circulation emerges throughout the embryo. The developing vasculature centers on cardiac function, forming a network intimately intertwined with neural architecture (**Fig. 4H, 4J, S9A, S9B, Movie S10**). Live imaging and growth factor expression confirm that neural development guides vasculogenesis (**Fig. 4I, 4J**). The circulatory system organizes into three distinct circuits within the brain, yolk sac, and spinal cord (**Fig. 4J, S9B, Movie S11**), with highly directional blood cell flow across the yolk sac surface (**Fig. 4J, S9B, Movie S11**). These findings support a model wherein the yolk sac facilitates organized mass transport beyond mere nutrient provision. Once established, blood flow speed remains homogeneous and constant throughout development (**Fig. 4K**). Pharmaceutical inhibition targeting angiogenesis (*31*) disrupts both neural and vascular development (**Fig. S9C**), confirming shared signaling pathways (*32, 33*).

To delineate the sequence of neurogenesis and organogenesis, we fixed transgenic embryos at distinct stages. These lines feature fluorescent labeling of vascular endothelium (**Fig. S1A**), neurons (**Fig. S1B**), lateral line (**Fig. S1C**), cardiomyocytes (**Fig. S1D**), skeletal muscle (**Fig. S1E**), and Ca²□ activity (**Fig. S1F**), with fluorescence onset marking process initiation. Concurrent phase-contrast imaging documented other organ formation. Sequential analysis (**Table S6**) verifies that neural development, coupled with histogenesis (Sox2, claudin, actin, myosin) (**Fig. 4A, 4B, S7, S8, S10A**), initiates prior to angiogenesis and organogenesis (**Fig. 2F, S10B-D**), and 30 hours before detectable neural activity (**Fig. 3J-P**). As a pervasive network, early neuronal architecture provides positional guidance for organ placement and vascular patterning, substantiating neurogenesis as the pioneering event in embryonic development.

## Summary and Discussion

In this work, we have elucidated zebrafish neurogenesis through a comprehensive *in vivo* imaging approach. Multi-channel imaging across temporal and spatial scales reveals the leading role of neurogenesis in early development, occurring concomitantly with histogenesis yet preceding organogenesis (**Fig. S11**). This distinctly orchestrated process unfolds through four sequential stages: somata clustering, network establishment, apoptosis-mediated optimization, and functional maturation. We have identified stereotypical phenomena during each stage, revealing potential early targets for brain health maintenance and Alzheimer’s disease intervention (*34*).

The neural system strategies revealed here provide fundamental insights into the developmental origins of intelligence and offer practical guidance for artificial intelligence design. Essentially, we can construct artificial systems using living organisms as blueprints. The observed pattern—where axons extend after somata form base-station-like clusters—ensures optimal mapping between neural architecture and body plan. The peak of apoptosis-mediated optimization immediately follows network establishment, when arbors remain simple, minimizing structural and metabolic costs. Its later persistence reflects a sophisticated system-level balance between information efficiency and energy expenditure. The delayed initiation of organ function after neural network establishment effectively prevents developmental dysfunction—a principle directly applicable to brain-inspired computing systems (*35*).

These findings raise critical questions for future investigation. First, the signaling pathways underlying the observed phenomena require molecular dissection at finer temporal resolution. Second, the material basis and functional significance of the slow transport phenomena demand clarification. Third, how network refinement and aging influence behaviors like swimming (*36*) merits exploration, particularly regarding neural-organ coordination. Finally, developing additional transgenic lines targeting early developmental regulators like Sox2 will further illuminate the coupling between stem cell differentiation and neurogenesis.

## Acknowledgments

We thank Tao Tan for helpful discussions; Peishi Wang and Mingyang Chen for helping maintain the transgenetic zebrafish lines; Runjie Yu and Siyi Zhang from Oujiang Laboratory for assistance in spinning disk confocal and two-photon microscopy imaging, respectively. This study was supported by taxpayers of China through the National Natural Science Foundation of China (distinguished young scholars funding, overseas) and Wuhan University (talents startup funding).

## Author contributions

Conceptualization: BL

Methodology: BL, ZDW

Investigation: BL, ZDW, LT

Visualization: ZDW

Funding acquisition: BL

Project administration: BL Supervision: BL

Writing – original draft: BL, ZDW

Writing – review & editing: BL, ZDW, LT

## Data and materials availability

All data are available in the main text or the supplementary materials.

## Supplementary Materials

### I. Materials and Methods

#### Construction and maintenance of transgenetic zebrafish lines

Zebrafish strains Tg(elavl3:GCaMP6f) and Tg(cldnb:lynEGFP) were acquired from the China Zebrafish Resource Center (http://www.zfish.cn/), whereas the Tg(elavl3:EGFP), Tg(cmlc2: EGFP), Tg(smyhc1: EGFP) and Tg(fli-1:mCherry) strains were purchased from Nanjing Ezerinka Biotechnology Co., Ltd. (http://www.ezerinka.com/). The hybrid lines (elavl3:EGFP; fli-1:mCherry) were generated in-house through crossbreeding and selective rearing. All zebrafish were maintained in a recirculating aquaculture system procured from Nanjing Ezerinka Biotechnology Co., Ltd.

The system temperature was set to 28□°C, with continuous operation of UV sterilization and oxygen supply. A 5% sea salt solution and 5% sodium bicarbonate solution were used to regulate conductivity and ion balance, maintaining system conductivity between 500–800 μS/cm and pH between 7.0–8.0. Ambient temperature was controlled at 28□°C using air conditioning, and a 12□h light/12□h dark photoperiod was applied (*1*). Approximately one-third of the system water was replaced daily to keep total ammonia nitrogen below 0.02□mg/L.

To facilitate subsequent spawning, male and female zebrafish were housed separately. Males were distinguished by their slender body shape, flat abdomen, and lemon-yellow coloration, whereas females exhibited a fuller body, rounded abdomen, and silvery-gray pigmentation, allowing reliable visual sex identification. Adult zebrafish were fed live Artemia nauplii daily. The hatching procedure was as follows: 5□g of Artemia cysts and 5□g of sea salt were added to 1□L of pure water. The mixture was aerated under illumination for 24□h, after which aeration was stopped to allow eggshells to separate. The hatched nauplii settled at the bottom and were collected for feeding. Fish were fed 1-2 times per day, with the amount adjusted to ensure consumption within 15-20 minutes.

#### Characterization of transgenetic zebrafish lines

The zebrafish strains used in this study are summarized in **Table S1**. Transgenic zebrafish lines were genotyped by examining the expression pattern of fluorescent proteins. Specifically, embryos at 48 hours post-fertilization (hpf) were observed under a fluorescence microscope, and the correct integration of the transgene was verified based on the specific localization of the fluorescent signal. For the identification of different transgenic zebrafish lines, we assessed the fluorescence expression in corresponding 48-hour-old embryos under a confocal microscope (**Fig. S1**), and assessed the key morphological features.

In the Tg(fli-1:mCherry) (*2*) line, the endothelial-specific promoter drives the expression of a red fluorescent protein, enabling visualization of the vascular system. As shown in **Fig. S1A**, distinct vascular-like structures were observed in the brain, heart, and spinal cord of zebrafish embryos. Blood cell flow within the vessels was also visible under bright-field microscopy, confirming correct labeling of this line. The Tg(elavl3:EGFP) (*3*) line utilizes a pan-neuronal-specific promoter fused with green fluorescent protein to visualize the nervous system. As demonstrated in **Fig. S1B**, strong EGFP expression was detected in the brain and spinal cord, with clearly visible neuronal somata and axonal networks, indicating proper labeling. In the Tg(cldnb:lynEGFP) (*4*) line, a lateral line system-specific promoter drives membrane-localized EGFP expression for visualization of the lateral line. **Fig. S1C** shows high EGFP signals in a rosette-like pattern at the lateral line positions along the spinal cord, confirming accurate labeling. The Tg(cmlc2:EGFP) (*5*) line expresses EGFP under a cardiomyocyte-specific promoter to visualize the heart. As observed in **Fig. S1D**, robust green fluorescence was present in the cardiac region, along with visible heartbeats, verifying correct labeling. In the Tg(smyhc1:EGFP) (*6*) line, a slow muscle-specific promoter drives EGFP expression for skeletal muscle visualization. **Fig. S1E** shows strong fluorescence in the spinal region with clearly defined muscle fibers, confirming proper labeling. The Tg(elavl3:GCaMP6f) (*7*) line uses a pan-neuronal promoter to express the calcium indicator GCaMP6f, enabling visualization of neural calcium dynamics. As shown in **Fig. S1F**, high fluorescence was observed in the brain and spinal cord, with discernible neuronal somata, axonal networks, and dynamic fluorescence fluctuations, indicating correct labeling.

#### Sample preparation for *in vivo* zebrafish imaging

On the day before imaging, female and male zebrafish of different transgenic strains were placed in spawning tanks separated by a divider. The tanks were filled with a 1:1 mixture of fresh water and system water, covered, and maintained overnight. On the following morning, after the onset of illumination, the divider was removed to allow direct interaction between males and females. Males began nudging the abdominal region of the females, which subsequently initiated spawning. Approximately 30 minutes later, both females and males were returned to the main rearing system. The fertilized eggs were collected from the spawning tanks into 10 cm Petri dishes and maintained in specialized zebrafish embryo medium at 28□°C (*1*).

Embryo imaging typically commenced around 24 hours post-fertilization (hpf). Prior to imaging, embryos were screened using a spinning disk confocal microscope, and only those exhibiting clear fluorescence signals were selected for further processing. The chorion was then removed from each embryo to improve imaging quality. Although the chorion serves as a protective barrier during early development, it limits imaging depth and introduces motion artifacts as embryos become more active inside the membrane. For dechorionation, 24-hpf embryos were transferred to a Petri dish with a small amount of water and placed under a stereomicroscope. A small incision was made in the chorion using a corneal scissors, and the embryo was gently extruded with fine forceps.

The dechorionated embryos were then transferred to a confocal imaging dish. Low-melting-point agarose (0.12□g) was dissolved in 10□mL of ddH2O by microwave heating to prepare a 1.2% solution (*8*). After cooling to approximately 35□°C, the agarose solution was dispensed over the embryos. The position and orientation of each embryo were adjusted with forceps before the agarose solidified. For imaging at earlier developmental stages, such as 12 hours post-fertilization (hpf), the chorion was not removed due to the structural fragility of the embryos at this period, as mechanical manipulation could readily cause damage. Instead, the embryos within their intact chorions were directly embedded in low-melting-point agarose for immobilization during imaging.

#### Live cell imaging

Live imaging of zebrafish embryos was performed using a Spin SR10 spinning disk confocal microscope (Evident Olympus) equipped with an Oko-lab incubation chamber (H301-MINI). The system maintained a constant temperature of 28□°C throughout the experiments. Embryos were mounted in confocal dishes (Nest, 801001) and imaged with a typical laser intensity set at 20% and exposure time at 200□ms. For fluorescence excitation, the 488□nm laser line was used to visualize pan-neuronal labels in the transgenic lines (elavl3:EGFP, cldnb:lynEGFP, elavl3:GCaMP6f, cmlc2:EGFP, and ssmyhc1: EGFP), while the 561□nm laser was employed for imaging the (fli-1:mCherry) line.

To enhance experimental throughput, approximately 20 embryos were placed in each confocal dish. The microscope’s positional memory function was utilized to record individual embryo locations, enabling simultaneous time-lapse acquisition from multiple specimens. All embryos were imaged using axial Z-stack acquisition mode with the following parameters. Z-stack imaging was performed with a 60x oil objective at 100 μm Z-range and 2-3 μm step size, a 30x oil objective at 150 μm Z-range and 2-3 μm step size, and a 10x objective at 400 μm Z-range and 10 μm step size, respectively. Time-series imaging was performed at 20-minute intervals over a total duration of 48 hours, generating continuous 3D volumetric data throughout the recording period (*9–11*).

The two-photon and light-field imaging were performed using FVMPE-RS (Evident Olympus) and LSM910 (Karl Zeiss), respectively. Two-photon imaging was performed using a 25× water immersion objective. With a z-step interval of 5 μm, the total imaging depth exceeded 500 μm. The stitching function was applied to achieve whole-embryo coverage of zebrafish, followed by three-dimensional (3D) reconstruction using the microscope’s proprietary software. For long-term volumetric imaging of zebrafish embryos with co-labeled neurons and blood vessels (elavl3:EGFP; fli-1:mCherry), the Light-sheet 4D Light-field Imaging Module of the LSM910 system was employed, with a 40× water immersion objective, a volumetric imaging depth of 110 μm, and a temporal imaging interval of 10 min.

#### Immunostaining Experiments

Zebrafish embryos at various developmental stages were collected after spawning and fixed with 4% paraformaldehyde (PFA; Pumeike, PMK0240) at room temperature for 2 hours. The chorions of the embryos were then carefully removed under a stereomicroscope. After repeated washing with PBS, the samples were permeabilized with 0.1% Triton X-100 (Solarbio, T8200-500ml) for 30 minutes at room temperature. Subsequently, blocking was performed using 3% bovine serum albumin (BSA; Pumeike, PMK0181-250g) containing 0.1% Triton X-100 for 2 hours at room temperature.

The embryos were incubated overnight at 4□°C in the dark with a SOX2 primary antibody (abcam, ab97959) diluted 1:500 in 3% BSA to label pluripotent stem cells and neural tem cells at early developmental stages (*12*). The following day, the antibody solution was removed, and the samples were washed three times with PBS (10 minutes per wash). Thereafter, the samples were incubated overnight at 4□°C in the dark with a mixture of Goat Anti-Rabbit IgG H&L (Alexa Fluor® 594; abcam, ab150080) (*13*) and Hoechst 33258 (*14*), both diluted 1:1000 in PBS. Finally, the samples were washed three times with PBS (10 minutes each), followed by two additional PBS washes, and then mounted on confocal dishes for imaging. The reagents used in the immunostaining and pharmaceutical interference experiments are summarized in **Table S2**.

#### Pharmaceutical interference experiment

The PI3K inhibitor LY294002 (*15, 16*) (MedChemExpress, HY-10108; supplied as a 10 mM solution in DMSO) was employed to investigate the PI3K/Akt signaling pathway. This pathway is a critical downstream component of both VEGF-driven angiogenesis, which promotes endothelial cell survival, proliferation, and migration, and neurotrophin signaling (e.g., NGF and BDNF), which is essential for neuronal survival and axonal growth. Accordingly, LY294002 disrupts these processes by inhibiting PI3K. For the experiments, a 1 mM stock solution was prepared by diluting the compound in DMSO. This stock was further diluted in zebrafish embryo medium to obtain working concentrations of 10, 50, and 100 µM. Zebrafish embryos were exposed to these solutions, and their neuronal and vascular development was assessed after 24 hours of treatment.

### II. Quantification and Analysis

#### Data reconstruction and 3D visualization

Whole-embryo imaging of the zebrafish neural network was performed using an Evident FVMPE-RS multiphoton laser scanning microscope. At 24 hours post-fertilization, zebrafish embryos (elavl3:EGFP) were dechorionated under a stereomicroscope and embedded in 1.2% low-melting-point agarose mounted at the bottom of a confocal dish, using either live or 4% PFA-fixed specimens. Image acquisition was carried out with a 25x water-immersion objective and a dedicated two-photon lens. Tiling was employed to cover the entire embryonic neural network, with Z-stacks acquired at 10 µm intervals, spanning a total depth of approximately 600 µm. The resulting datasets were processed using Olympus imaging software for three-dimensional reconstruction. Alternatively, Z-stack images were imported into Imaris (Oxford Instruments) for further 3D rendering and visualization.

The reconstruction of imaging stacks from confocal microscopy is conducted using Cell Sens Dimension Desktop 4.2.1 (https://lifescience.evidentscientific.com.cn/en/software/cellsens/). In the top planar view, the Maximum Intensity Projection (MIP) algorithm is adopted to overlay all images in the z-stack into a single image. In the side view, the Marching Cubes (Surface Reconstruction Algorithms) algorithm is used to yield the stereoscopic spheroids (**Fig. S3A**). Compared with the range in the xy plane, the length scale in z is relatively small, making the analysis based on xy projection reasonably accurate.

#### Analysis of live cell imaging data

Time-lapse imaging data of zebrafish neural network development were analyzed using Fiji (ImageJ). Growth cone dynamics, including migration trajectories and velocity, were quantified with the Manual Tracking plugin (*17*). The analysis workflow consisted of the following steps: Time-lapse image stacks were loaded into Fiji, and the Manual Tracking plugin was initialized with appropriate temporal intervals and spatial calibration based on the acquisition settings. Growth cones were manually selected and tracked across consecutive frames using the plugin’s point-and-click interface. The plugin automatically recorded their positional coordinates over time and computed kinematic parameters such as displacement, velocity, and persistence. To assess structural organization within the developing network, angular relationships between neurites were also measured using Fiji’s built-in angle measurement tool. Specific neurite segments and growth cone orientations were manually annotated, and their relative angles were calculated to evaluate network topology and directional guidance. All tracking and angular data were exported for further statistical analysis and graphical representation in GraphPad Prism. The average fractal dimension (DC) was used to quantify the branching complexity of neural dendrites via the box-counting method in the FracLac plugin (v.2016Apr1) for ImageJ (v.1.8.0_172): neuronal images were converted to 8-bit grayscale, binarized with the Otsu algorithm (foreground preserved as native morphology), and analyzed with parameters including 6 multi-grid scan positions, Powered Series scaling (base=2, sampling sizes 2-64 pixels, n=4), locked background color, and enabled “slip grid” and “check pix” options; DC was calculated as the mean of grid-derived fractal dimensions (each derived from the negative slope of ln(box count) vs. ln(1/sampling size) regression, r²>0.99, σ<0.02 for reliability).

#### Transcriptome sequencing and analysis

Zebrafish embryos were collected at 12, 24, 36, 48, 60, and 72 hours post-fertilization (hpf). Approximately ten embryos per time point were immediately placed in RNase-free tubes containing TRIzol reagent and subsequently sent to GeneChem (Shanghai, China) for RNA sequencing. The resulting transcriptomic data were analyzed in our laboratory.

Gene set enrichment analysis was carried out using R software (v4.2.1) with the cluster Profiler package. Neural-associated highly expressed genes from distinct developmental windows were analyzed for KEGG pathway enrichment (*18*). Significantly enriched terms were identified across multiple functional categories, including Biological Process (BP), Molecular Function (MF), and Cellular Component (CC), with a focus on pathways related to neural development, synaptic transmission, and axon guidance. Visualization of temporal expression trends and functional enrichment results was conducted using ggplot2 (*19*) in R, enabling clear representation of dynamic transcriptome changes during embryonic neural development.

The resulting gene expression matrices across time points were subjected to subsequent bioinformatic analysis (**Fig. S3B, S3C**). Time-series gene clustering was performed using the online platform bioinformatics.com.cn with the Mfuzz package, which identifies genes with similar expression dynamics throughout development. Clusters exhibiting neural-related expression patterns were selected for further functional annotation.

### III. Additional Discussion

#### Interplay among Sox2 expression, MVN neurogenesis, and lateral line development

During zebrafish embryonic development, the transcription factor Sox2 shows a highly dynamic and functionally specific spatiotemporal distribution, and its transition from early embryonic stem cells to neural stem cells is a core feature of the conserved mechanism underlying vertebrate neural development. Existing evidence clearly indicates that Sox2 is expressed in pluripotent cell populations during early zebrafish embryogenesis and becomes specifically enriched in neural stem cells (NSCs) or neural progenitor cells (*20, 21*) as the nervous system initiates development, a process that is critical for neural plate formation, neural tube construction, and subsequent neuronal differentiation. Immunofluorescence staining of Sox2 at different developmental stages reveals that at the cleavage stage (3 hpf), all cells highly express Sox2 (**Fig. S7A**), at 12 hpf, cells at the animal pole exhibit strong Sox2 expression (**Fig. S7B**), at 24 hpf, Sox2 is highly expressed in neural stem cells, especially in the brain region, with two regularly arranged columns of Sox2□high cells present along the spinal cord (**Fig. 4A, S7C**), at 36 hpf, the Sox2 expression pattern is similar to that at 24 hpf, but Sox2 expression in the spinal cord tends to shift toward the location of the MVN and posterior lateral line (**Fig. S7D**), at 48 hpf, the Sox2 expression pattern differs significantly from that at 36 hpf, with only one highly organized column of Sox2□high cells remaining in the spinal cord that spatially overlaps with the MVN (**Fig. 4B, S7E**). Immunofluorescence staining targeting the lateral line further demonstrates that the posterior lateral line spatially co□localizes with Sox2 at 48 hpf (**Fig. 4C, S8B**), and time□lapse imaging of fluorescently labeled lateral line zebrafish embryos shows that the posterior lateral line forms approximately from 34 hpf to 48 hpf, while the MVN forms from around 27 hpf to 38 hpf (**Fig. 4C**). Based on the developmental timeline and spatial localization, we conclude that during neural development, Sox2□high cells are consistently positioned earlier at the sites where subsequent neurogenesis takes place, the MVN likely plays a leading role in posterior lateral line development, and the posterior lateral line may develop along the pre□established MVN pathway.

#### Gene co-expression analysis

To further explore the relationships between neurogenesis and stem cells, cell death, and calcium signaling, we performed additional analyses of the transcriptomic data at different developmental stages. We conducted time-series analysis of neural-related, stem cell–related, cell death–related, and calcium-related genes in zebrafish embryos at distinct developmental time points, identified highly expressed genes at each stage, and then used Venn diagrams to visualize genes that were simultaneously upregulated across different biological processes.

The key genes that are involved in both neural processes and cell death during zebrafish embryonic development are summarized in **Table S3**. By screening highly expressed genes at each stage and constructing a Venn diagram (**Fig. 3C**), we identified shared differentially expressed genes between the two groups. Analysis of these co-expressed genes reveals that neural network development and apoptotic pruning share key regulatory genes, demonstrating that neural development and programmed cell death are highly coordinated rather than independent processes. These shared genes participate in axon formation and neural network assembly, as well as in the elimination of abnormal or redundant neurons.

The key genes that are involved in both neural processes and calcium signaling during zebrafish embryonic development are summarized in **Table S4**. By screening highly expressed genes at each stage and constructing a Venn diagram (**Fig. 3C**), we identified shared differentially expressed genes between the two groups. Analysis of these co-expressed genes reveals that neural development is highly coupled with calcium signaling pathways, and calcium signaling serves as a core regulatory mechanism for early zebrafish neurogenesis. These shared genes are simultaneously involved in neural differentiation, axon growth, synaptogenesis, and neural network maturation, and directly regulate calcium influx, calcium transients, calcium wave propagation, and calcium homeostasis.

The key genes that are involved in both neural processes and stem cell differentiation during zebrafish embryonic development are summarized in **Table S5**. By screening highly expressed genes at each stage and constructing a Venn diagram (**Fig. 4F**), we identified co-upregulated differentially expressed genes shared by the two groups. Analysis of these co-expressed genes reveals that neurogenesis is tightly coupled with stem cell pluripotency regulation. During early zebrafish neural development, neural differentiation and stem cell pluripotency maintenance or exit share a core set of regulatory genes, indicating that neurogenesis is not simply the initiation of differentiation but proceeds synchronously and cooperatively with stem cell state transition. The core shared genes control both neural differentiation and the stem cell proliferation–differentiation switch. These genes exert dual functions: on the one hand, they promote neuronal maturation, axon growth, and neural network assembly; on the other hand, they suppress excessive stem cell proliferation, drive cell cycle exit, and determine the fate of neural progenitor cells.

#### Pharmaceutical interference experiment

The chemical compounds, antibodies, and live-cell dyes employed are listed in Table 2. LY294002 is a broad-spectrum PI3K inhibitor targeting PI3Kα, PI3Kδ, and PI3Kβ with IC50 values of 0.5, 0.57, and 0.97 μM, respectively (*15, 22, 23*). It also inhibits casein kinase 2 (CK2) and acts as a competitive DNA-PK inhibitor, binding reversibly to its kinase domain with an IC50 of 1.4 μM (*22*). As an apoptosis inducer (*16*), LY294002 impairs neural (*24*) and vascular development primarily through suppression of the PI3K/Akt signaling pathway. In neural development, it attenuates the anti-apoptotic activity of Akt, leading to increased neuronal cell death, particularly among embryonic neural precursors (*24*). Furthermore, it disrupts neuronal migration (*25*) and differentiation (*26*), resulting in mislocalization and altered cell fate specification, which ultimately compromises neural circuit assembly. Axonal outgrowth and synaptogenesis are also suppressed due to impaired cytoskeletal reorganization and growth cone dynamics (*27, 28*). In vascular development, LY294002 inhibits (*29*) by blocking VEGF-induced Akt activation, thereby reducing angiogenic output (*30*). It also interferes with EC migration (*30*) and vascular branching (*31*) by dampening chemotactic responses and motility, and undermines vessel stability by reducing pericyte coverage, leading to fragile and dysfunctional vasculature. For the experiments, a 1 mM stock solution was prepared by diluting the compound in DMSO.

This stock was further diluted in zebrafish embryo medium to obtain working concentrations of 10, 50, and 100 µM. Zebrafish embryos were exposed to these solutions, and their neuronal and vascular development was assessed after 24 hours of treatment. Consistent with these mechanisms, our experimental data show that treatment of zebrafish embryos with 100 μM LY294002 for 24 hours significantly elevated embryonic lethality and delayed both neural and vascular development.

#### Sequence of Developmental Events in Zebrafish Embryogenesis

Based on long-term live-cell imaging of different transgenic zebrafish lines and immunofluorescence staining analyses of the stem cell marker gene sox2 at various developmental stages, we summarized the temporal order of key developmental events in zebrafish embryos. The initial timing of each event during zebrafish embryogenesis is summarized in **Table S6**.

As a stem cell transcription factor, sox2 is expressed earliest. Strong sox2 signals were detected as early as 3 hours post-fertilization (hpf) in zebrafish embryos, with expression observed in all cells at the animal pole. By 24 hpf, sox2 was highly expressed in neural stem cells.

In the transgenic zebrafish line Tg(smyhc1:EGFP), the specific promoter smyhc1 (slow myosin heavy chain 1) encodes the myosin heavy chain protein essential for slow muscle contraction, regulates the ordered assembly of myomeres and the normal contractile function of slow muscle (*6*), and is specifically expressed only in slow skeletal muscle. Using live-cell imaging, the relevant signal was detected at approximately 15 hpf.

In the transgenic zebrafish line Tg(elavl3:EGFP), elavl3 belongs to the ELAVL family of RNA-binding protein genes. Its core function is to regulate gene expression at the post-transcriptional level and stabilize key mRNAs that maintain neuronal identity and function; it is specifically expressed only in neurons (*3*). Its promoter contains neuron-specific cis-regulatory elements that drive early, specific, and widespread expression of downstream reporter genes, making it a classic tool for pan-neuronal live imaging in zebrafish. The relevant signal was detected at approximately 18 hpf via live-cell imaging.

In the transgenic zebrafish line Tg(fli-1:mCherry), fli-1 is a key transcription factor gene of the ETS family. Its core functions include regulating the differentiation, proliferation, and migration of vascular endothelial cells during embryonic development, and mediating angiogenesis and hematopoietic development (*2*). It is specifically expressed mainly in vascular endothelial cells and some hematopoietic cells. Its promoter is highly vascular-specific and can efficiently drive downstream gene expression, serving as a common tool for vascular visualization studies in zebrafish. The relevant signal was detected at approximately 22 hpf via live-cell imaging.

In the transgenic zebrafish line Tg(cmlc2:EGFP), cmlc2 (cardiac myosin light chain 2) belongs to the myosin light chain family and is a marker gene of zebrafish myocardial tissue (*5*). Its core function is to encode the myosin light chain protein essential for cardiac contraction, participate in the assembly of cardiac myomeres, and regulate the contraction and relaxation of myocardial fibers, thereby maintaining normal cardiac rhythm and pumping function. It is specifically expressed only in cardiomyocytes. The relevant signal was detected at approximately 22 hpf via live-cell imaging, coinciding with the timing of vascular development, which may be attributed to the coordinated function of blood vessels and the heart.

In the transgenic zebrafish line Tg(cldnb:lynEGFP), cldnb (claudin b) belongs to the claudin family of tight junction proteins. Its core functions include participating in the assembly and maintenance of intercellular tight junctions, regulating cell polarity, cell migration, and tissue barrier formation (*4*). It is specifically and highly expressed in lateral line-related structures such as the posterior lateral line primordium and neuromasts in zebrafish. As a tight junction protein, the corresponding fluorescent signal can be observed on the embryonic surface before 24 hpf; however, lateral line-specific signals become detectable at approximately 35 hpf.

Based on the above experimental data and current progress in developmental biology, early zebrafish development follows a precisely orchestrated sequence: from stem cell specification to early histogenesis, primary organogenesis, and finally to sensory and functional maturation. The ubiquitous expression of sox2 at 3 hpf represents the establishment of developmental potency. Before germ layer differentiation, it is essential to maintain a highly proliferative and undifferentiated cell pool. By 24 hpf, the restricted expression of sox2 in neural stem cells marks the transition from broad pluripotency to the maintenance of lineage□specific progenitor cells, directly providing the cellular source for subsequent neurogenesis.

Complex organ assembly relies on the prior formation of tissues (histogenesis). The early emergence of the cytoskeleton components, including actin (3 hpf, SiR actin), myosin (15 hpf, smyhc1), claudin (24 hpf, cldnb), establishes the fundamental mechanical and structural axes of the embryo. The intertwined initiation of neuronal differentiation (18 hpf, elavl3) indicates that basic sensory□motor pathways are pre□patterned before the formation of visceral organ systems. A key insight from the timeline is that distinct neurogenesis (∼18 hpf) precedes the formation of vascular and cardiac networks (∼22 hpf). During early embryonic development, the nervous system often acts as a spatiotemporal template. The early□established neural axons and glial structures secrete essential guidance cues and form physical pathways that induce subsequent organogenesis. Notably, the early neural network may pre□pattern the trajectories subsequently followed by vascular endothelial cells (fli□1), ensuring that the development of the circulatory system precisely matches the high metabolic demands of the nervous system.

Organogenesis requires precise synchronization among multiple tissue types. The simultaneous appearance of vascular (fli□1) and cardiac (cmlc2) signals at 22 hpf provides a clear example. A functional circulatory system cannot form without the coordination of the heart and blood vessels, which are therefore tightly coupled via shared regulatory pathways. Once these life□sustaining primary organs are established, the embryo can support the histogenesis of more delayed, specialized secondary structures, such as the migration and assembly of the lateral line system (35 hpf, cldnb).

---

**Fig. S1.**
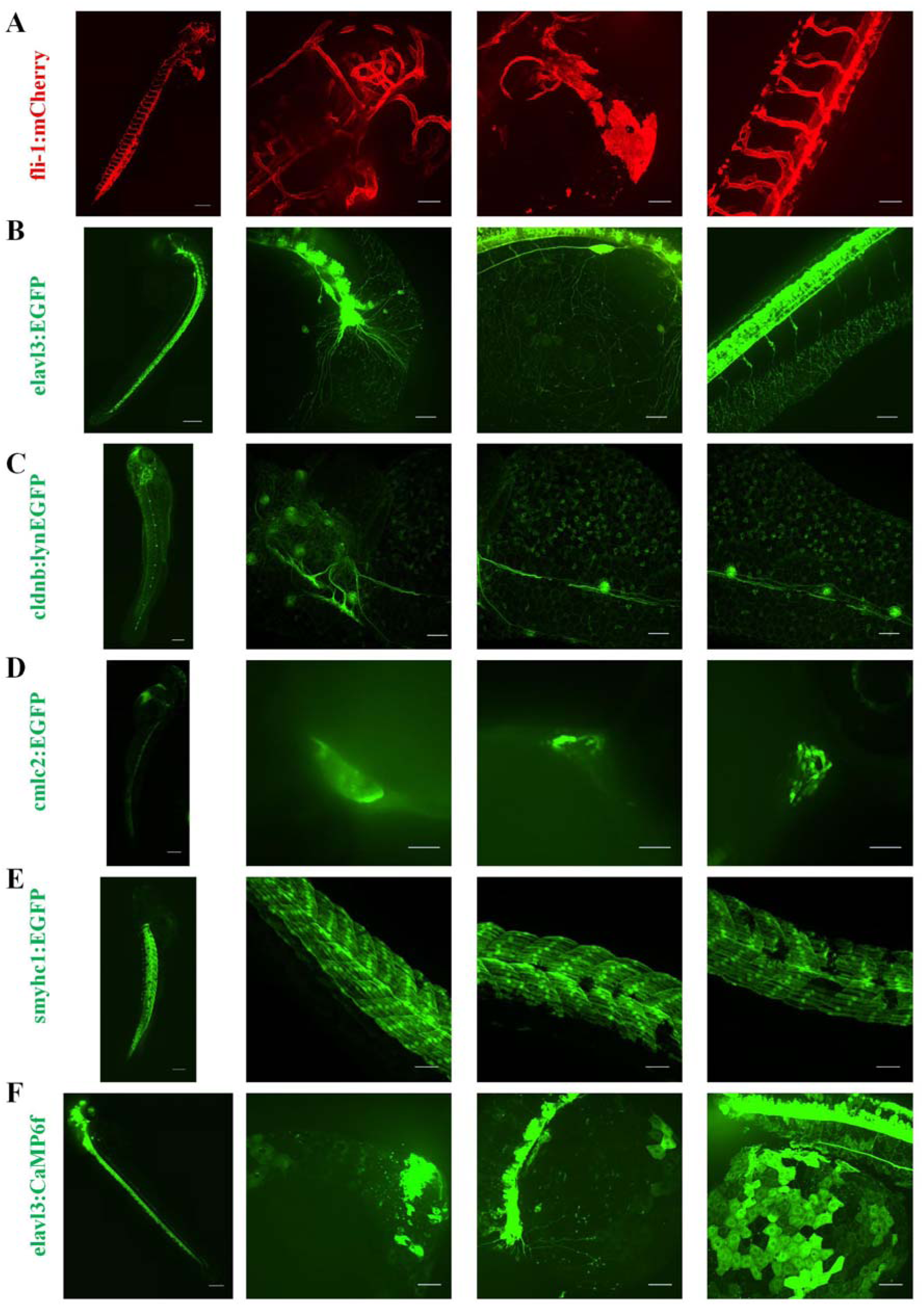
The fluorescently labeled transgenetic zebrafish line. (**A**) Representative confocal image of a zebrafish (fli-1:mCherry) embryo. Red fluorescence labels vascular endothelium. The left panel shows the entire embryo; scale bar = 200 μm. The right panel shows a magnified view; scale bar = 50 μm. (**B**) Representative confocal image of a zebrafish (elavl3:EGFP) embryo. Green fluorescence labels pan-neuronal structures. The left panel shows the entire embryo; scale bar = 200 μm. The right panel shows a magnified view; scale bar = 50 μm. (**C**) Representative confocal image of a zebrafish (cldnb:lynEGFP) embryo. Green fluorescence labels the lateral line system. The left panel shows the entire embryo; scale bar = 200 μm. The right panel shows a magnified view; scale bar = 50 μm. (**D**) Representative confocal image of a zebrafish (cmlc2:EGFP) embryo. Green fluorescence labels cardiac muscle. The left panel shows the entire embryo; scale bar = 200 μm. The right panel shows a magnified view; scale bar = 50 μm. (**E**) Representative confocal image of a zebrafish (smyhc1:EGFP) embryo. Green fluorescence labels skeletal muscle. The left panel shows the entire embryo; scale bar = 200 μm. The right panel shows a magnified view; scale bar = 50 μm. (**F**) Representative confocal image of a zebrafish (elavl3:CaMP6f) embryo. Green fluorescence reflects neuronal calcium activity. The left panel shows the entire embryo; scale bar = 200 μm. The right panel shows a magnified view; scale bar = 50 μm.

**Fig. S2.**
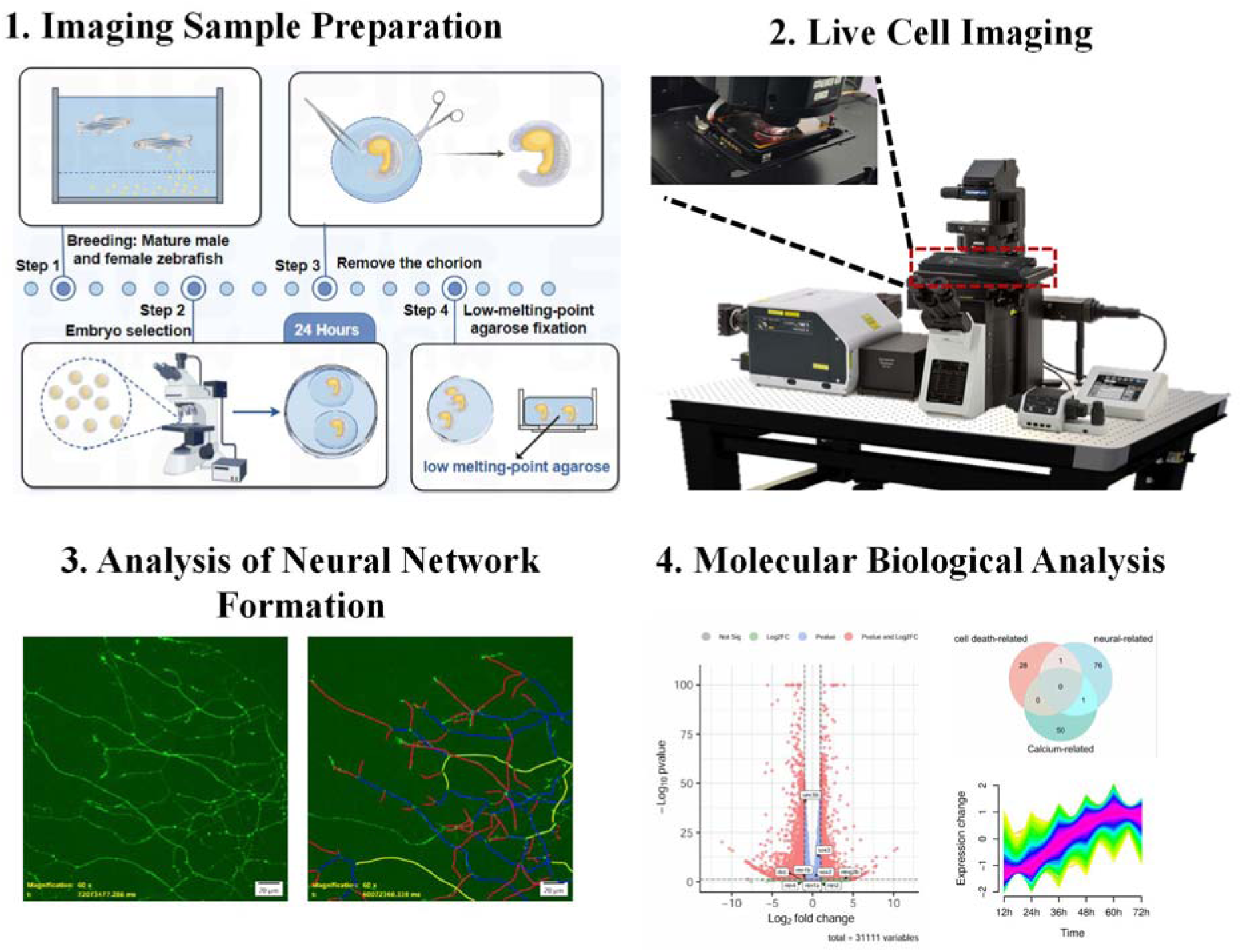
Schematics of the experimental design. An integrated approach of live imaging and transcriptomics was employed. First, zebrafish embryos were prepared by dechorionation and mounting in agarose for stable long-term imaging under controlled environmental conditions. Following this, time-lapse image data were acquired and analyzed. Concurrently, transcriptome sequencing was performed on embryos collected at different developmental stages. Finally, the imaging dynamics and gene expression data were correlated to identify potential functional links.

**Fig. S3.**
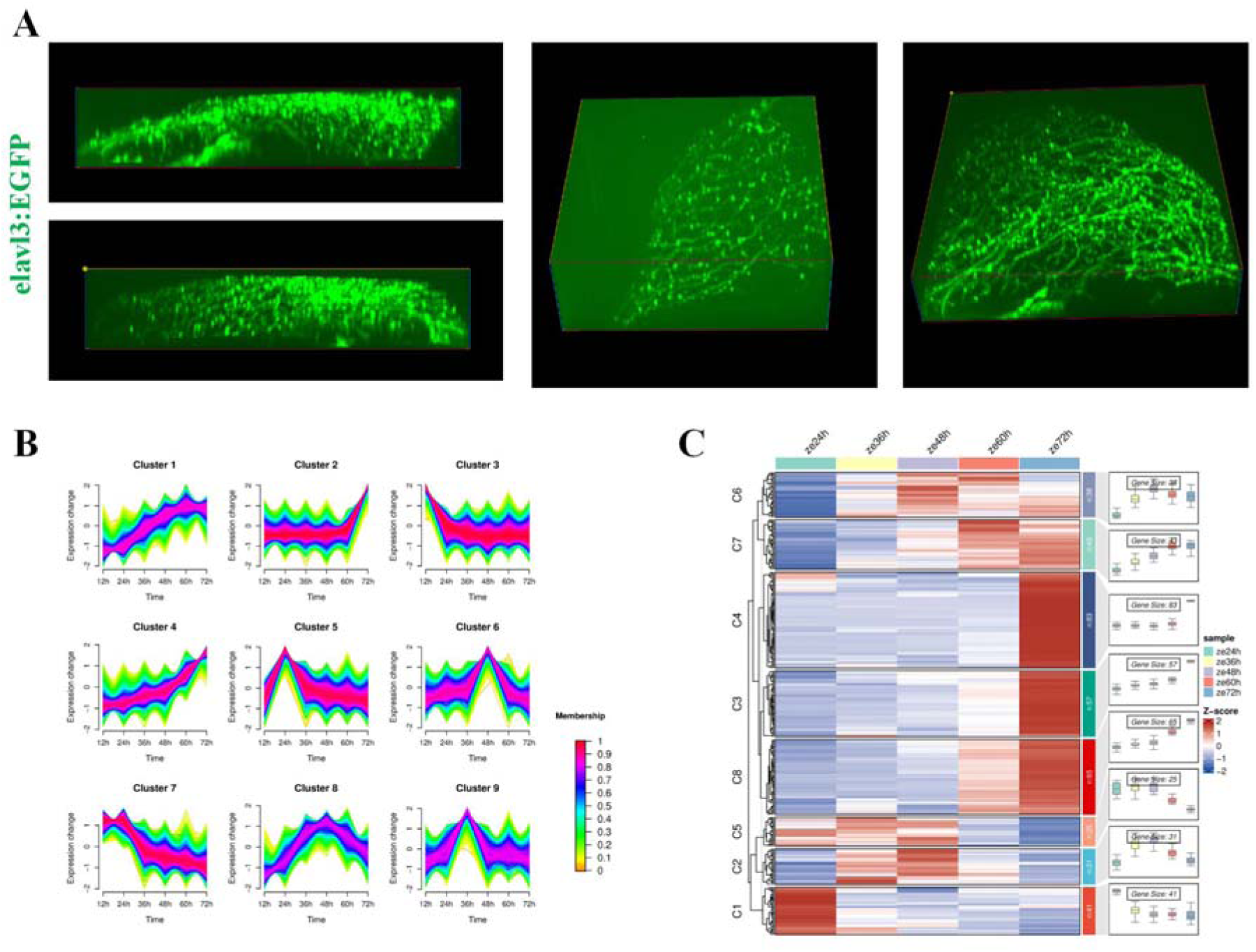
Imaging reconstruction and transcriptomic analysis of zebrafish embryos. (**A**) Representative image of a three-dimensional reconstruction of the embryonic neural network in zebrafish, generated using CellSens Dimension Desktop software (version 4.2.1). (**B**) Genome-wide Mfuzz time-series gene clustering analysis of zebrafish embryos across various developmental stages. (**C**) Heatmap and corresponding expression trend profiles of genome-wide transcriptome dynamics during zebrafish embryonic development. micrographs of a Tg(elavl3:EGFP) zebrafish embryo during the initial phase of neurogenesis, acquired at a high temporal resolution (20-minute intervals). EGFP signal (green) labels neurons. Scale bar= 200 μm. (**B**) Time-lapse confocal imaging (20-minute intervals) of a Tg(elavl3:EGFP) embryo, showing the extension of neural networks from cerebral neuronal somata toward the brain, spinal cord, and yolk sac. EGFP signal (green) marks neurons. Scale bar=50 μm. (**C**) Representative high-temporal-resolution confocal images (20-minute intervals) capturing the caudal growth of the primary ventral nerve toward the spinal cord in a Tg(elavl3:EGFP) embryo during neural development. Neurons are visualized by EGFP fluorescence (green). Scale bar= 100 μm.

**Fig. S4.**
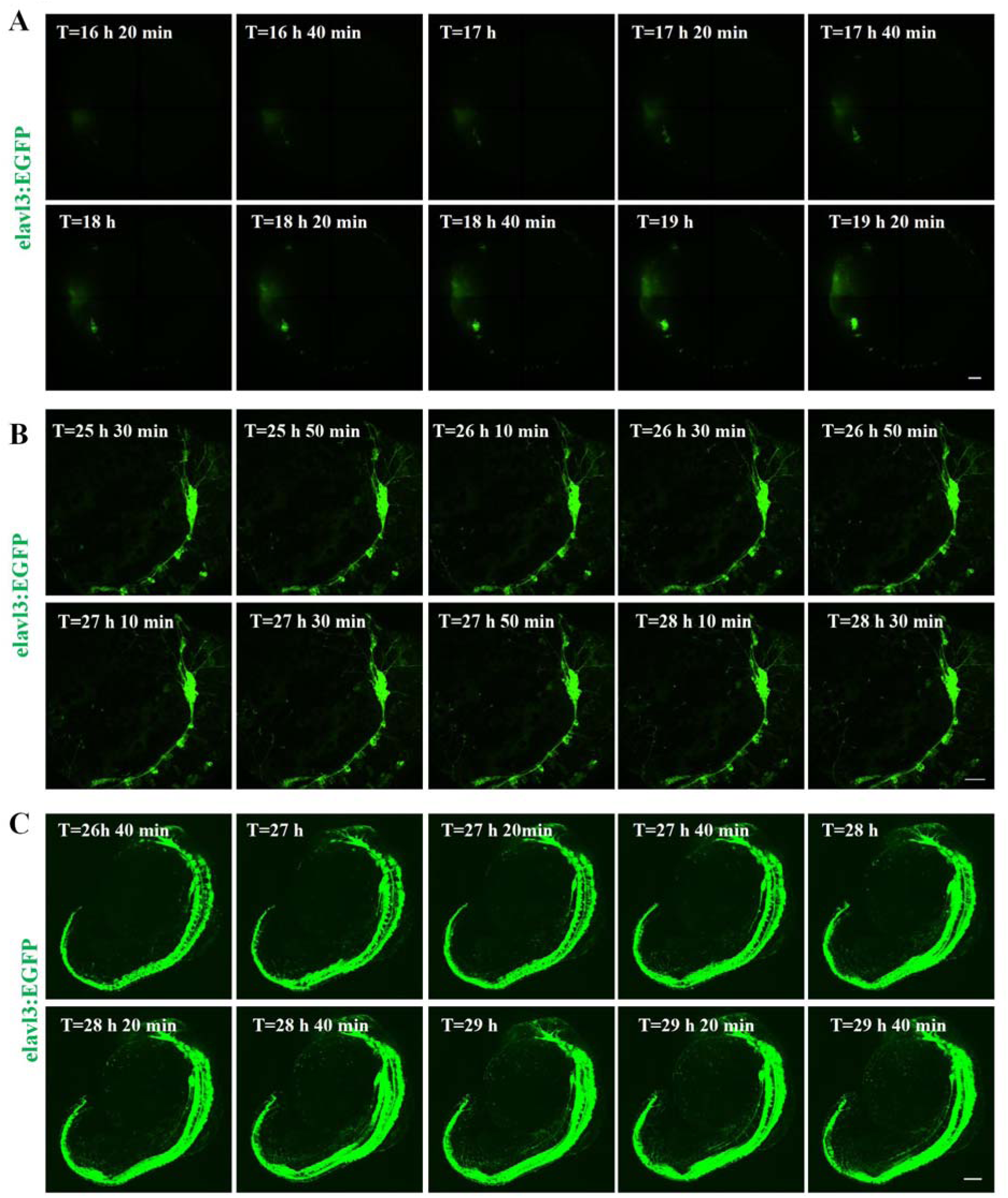
Raw images of initial stages at a finer time scale. (**A**) Time-series confocal micrographs of a Tg(elavl3:EGFP) zebrafish embryo during the initial phase of neurogenesis, acquired at a high temporal resolution (20-minute intervals). EGFP signal (green) labels neurons. Scale bar= 200 μm. (**B**) Time-lapse confocal imaging (20-minute intervals) of a Tg(elavl3:EGFP) embryo, showing the extension of neural networks from cerebral neuronal somata toward the brain, spinal cord, and yolk sac. EGFP signal (green) marks neurons. Scale bar=50 μm. (**C**) Representative high-temporal-resolution confocal images (20-minute intervals) capturing the caudal growth of the primary ventral nerve toward the spinal cord in a Tg(elavl3:EGFP) embryo during neural development. Neurons are visualized by EGFP fluorescence (green). Scale bar= 100 μm.

**Fig. S5.**
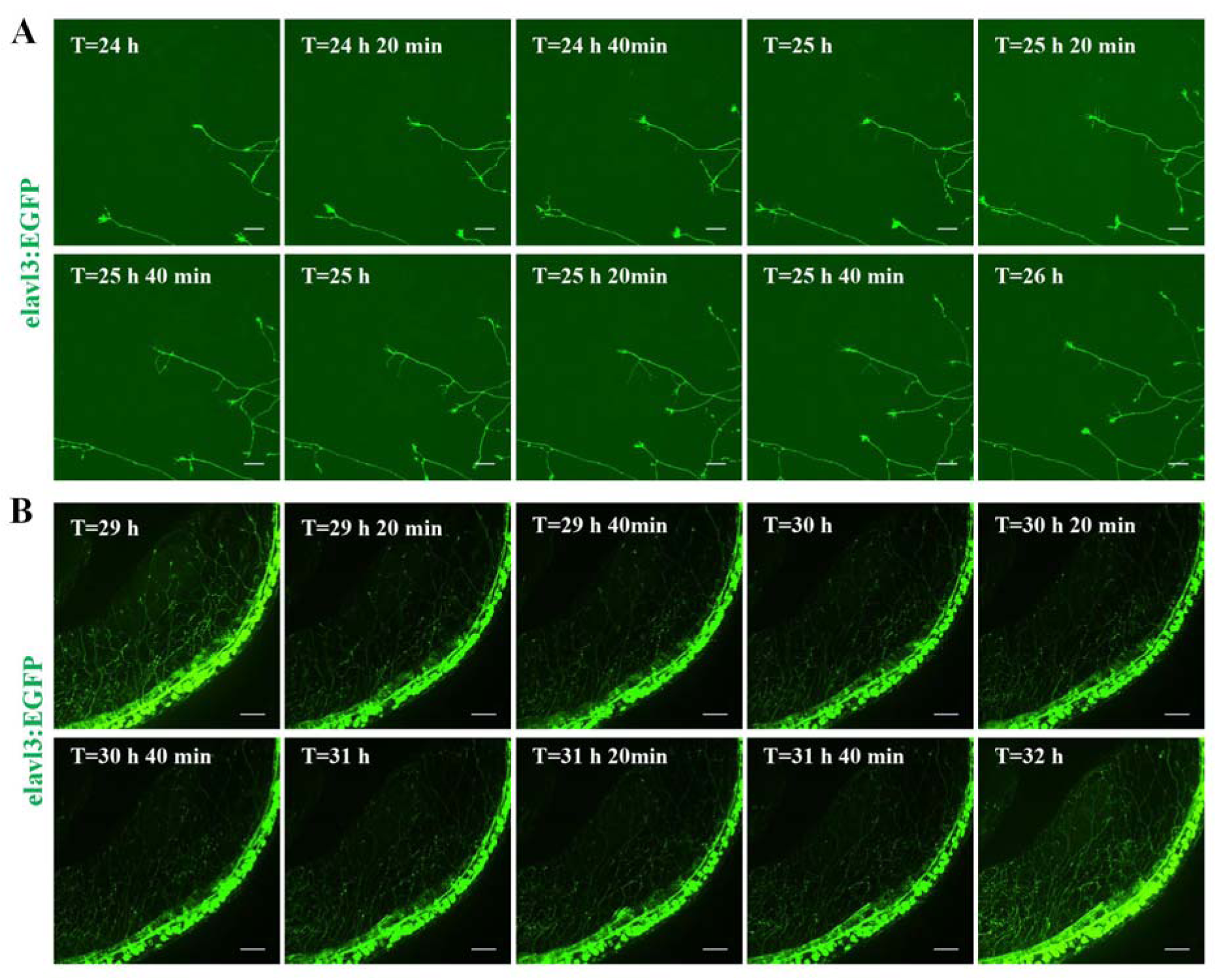
Raw images of the network formation in both brain and spinal cord at a finer time scale. (**A**) Representative confocal images showing the dynamic process of neural network formation on the zebrafish brain and yolk sac surface at higher temporal resolution. The time interval between the two images is 20 minutes. Scale bar = 20 μm. (**B**) Representative confocal images illustrating the dynamic process of neural network formation in the zebrafish spinal cord at higher temporal resolution. The time interval between the two images is 20 minutes. Scale bar = 50 μm. time-series confocal images showing partial neural networks undergoing apoptosis during neural development in zebrafish. White arrows indicate neural networks undergoing apoptosis. Scale bars: 50 μm in A and C, 20 μm in B. (**D and E**) Representative confocal images of cargo transport in the mature neural network of transgenic zebrafish embryos. Panels show the D elavl3:EGFP and E elavl3:GCaMP6f lines. For each, the left image provides a low-magnification overview. The scale bar is 50 µm, and the right image is a higher-magnification view of the boxed region, with transported cargo indicated by white arrows. The scale bar is 10 µm.

**Fig. S6.**
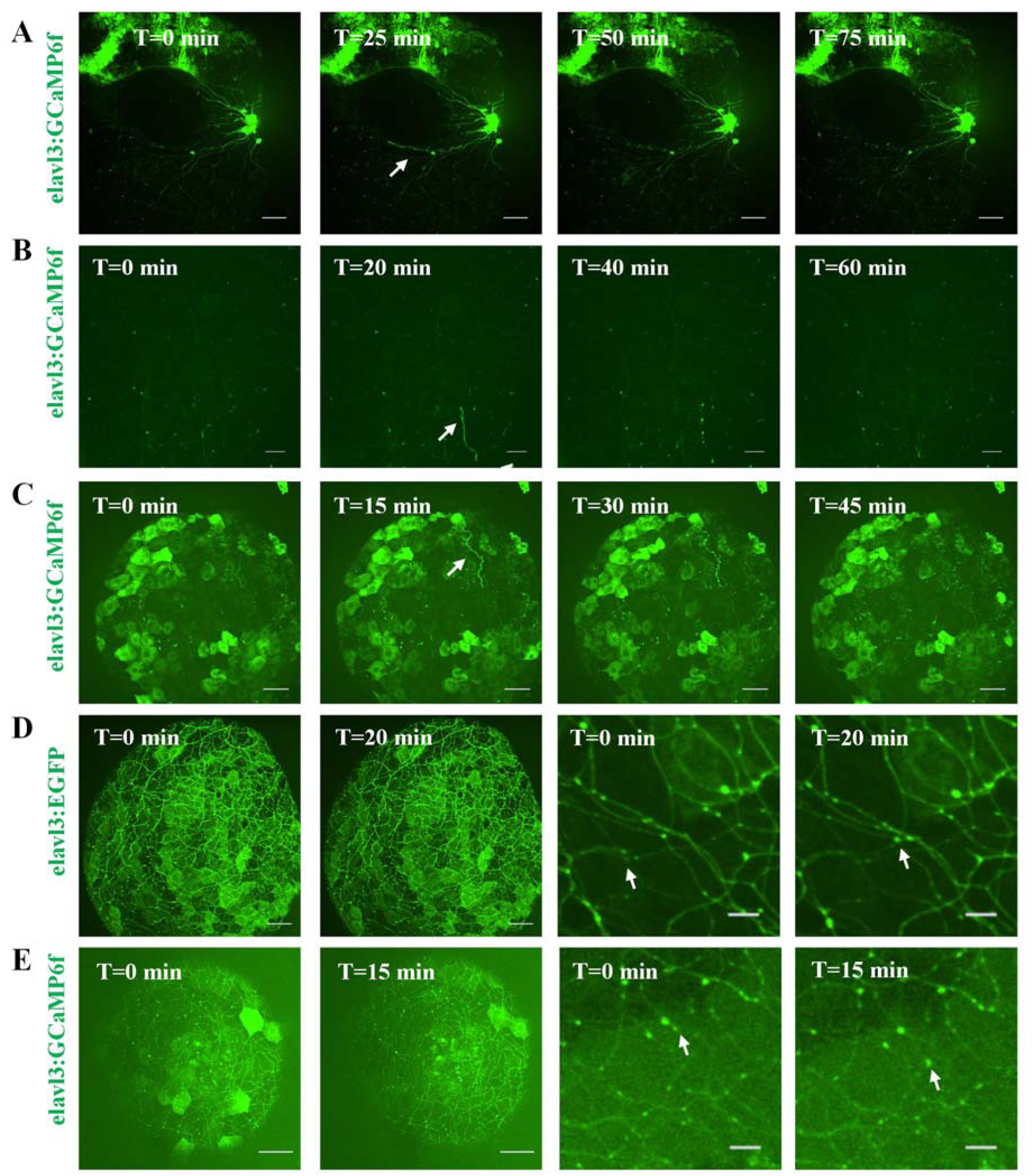
More examples of apoptosis and brain functions. (A, B and. **C)** Representative time-series confocal images showing partial neural networks undergoing apoptosis during neural development in zebrafish. White arrows indicate neural networks undergoing apoptosis. Scale bars: 50 μm in A and C, 20 μm in B. (**D and E**) Representative confocal images of cargo transport in the mature neural network of transgenic zebrafish embryos. Panels show the D elavl3:EGFP and E elavl3:GCaMP6f lines. For each, the left image provides a low-magnification overview. The scale bar is 50 µm, and the right image is a higher-magnification view of the boxed region, with transported cargo indicated by white arrows. The scale bar is 10 µm.

**Fig. S7.**
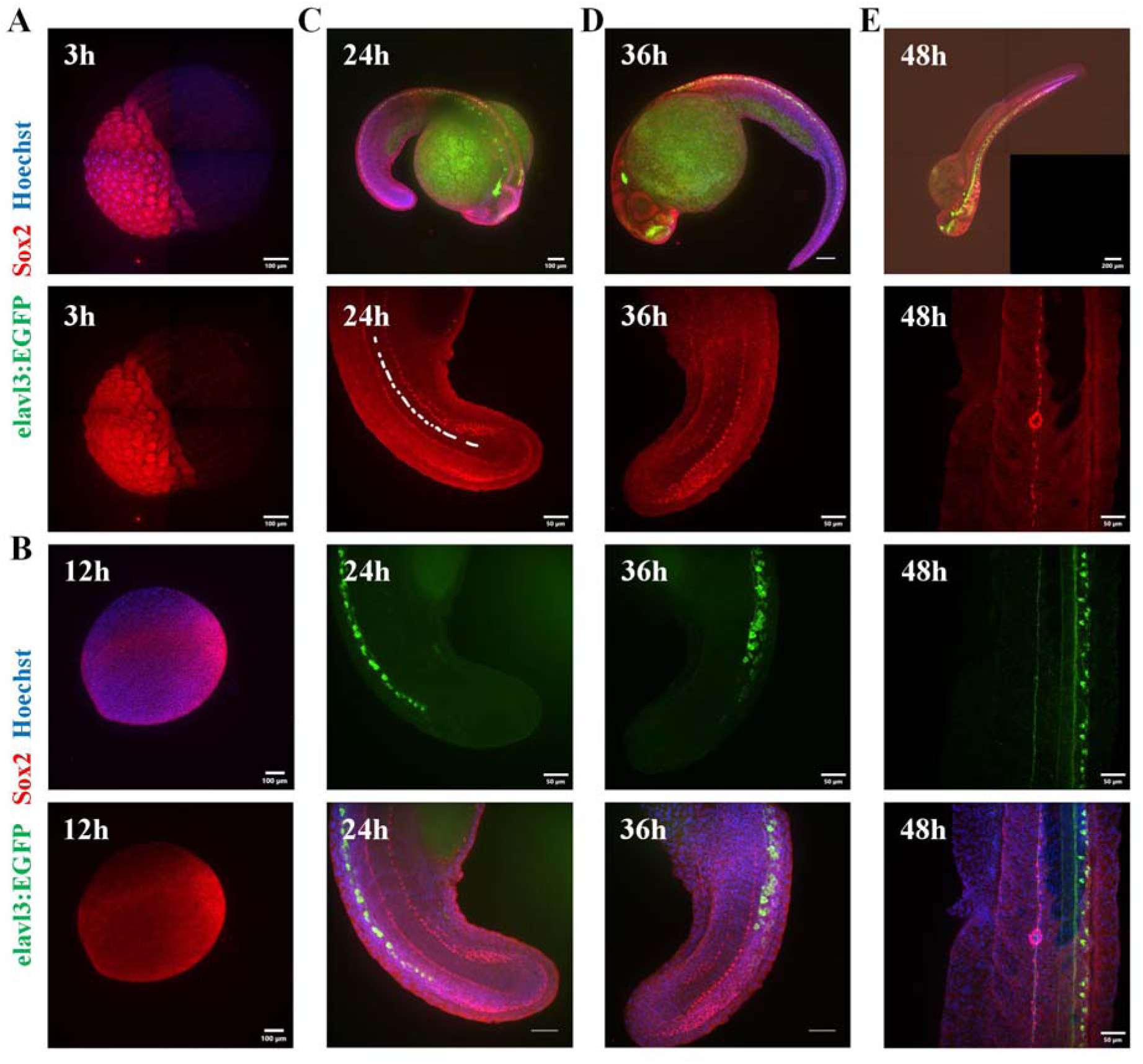
Correlation between sox2 and neural development during zebrafish embryogenesis. (**A**) Immunofluorescence staining of Sox2 in zebrafish embryos at 3 hpf. Sox2 was labeled with red fluorescence. Scale bar = 100 μm. (**B**) Immunofluorescence staining of Sox2 in zebrafish embryos at 12 hpf. Sox2 was labeled with red fluorescence. Scale bar = 100 μm. (**C**) Immunofluorescence staining of Sox2 and neurons in Tg(elavl3:EGFP) zebrafish embryos at 24 hpf. Sox2 was labeled with red fluorescence, and pan-neurons were labeled with green fluorescence. The upper panel was captured with a 10× objective, Scale bar = 100 μm; the lower panel was captured with a 30× silicone oil objective, Scale bar = 50 μm. (**D**) Immunofluorescence staining of Sox2 and neurons in Tg(elavl3:EGFP) zebrafish embryos at 36 hpf. Sox2 was labeled with red fluorescence, and pan-neurons were labeled with green fluorescence. The upper panel was captured with a 10× objective, Scale bar = 100 μm; the lower panel was captured with a 30× silicone oil objective, Scale bar = 50 μm. (**E**) Immunofluorescence staining of Sox2 and neurons in Tg(elavl3:EGFP) zebrafish embryos at 48 hpf. Sox2 was labeled with red fluorescence, and pan-neurons were labeled with green fluorescence. The upper panel was captured with a 10× objective, Scale bar = 100 μm; the lower panel was captured with a 30× silicone oil objective, Scale bar = 50 μm.

**Fig. S8.**
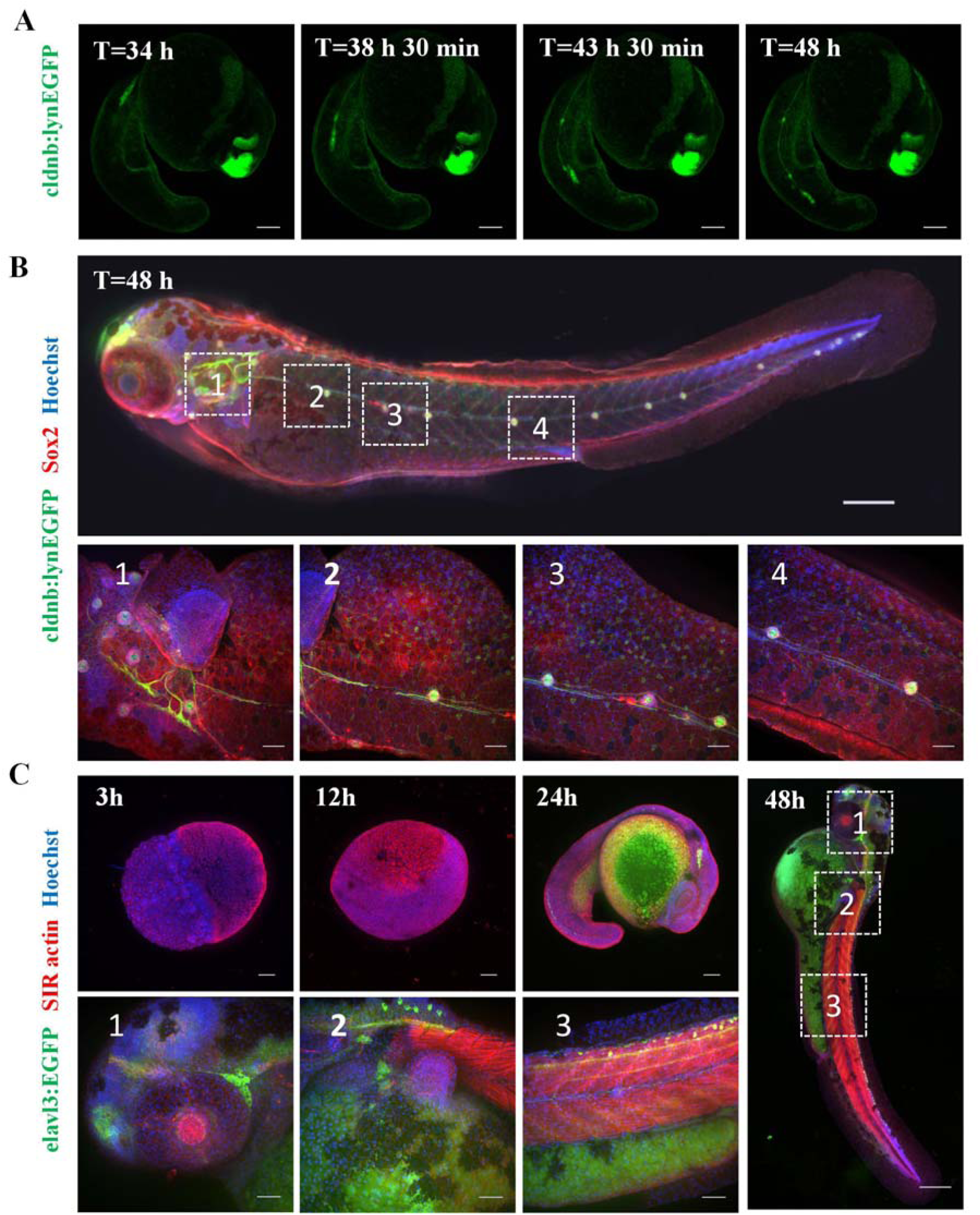
Relationship between neurogenesis and organogenesis. (**A**) Representative confocal images of posterior lateral line formation during zebrafish (cldnb:lynEGFP) embryonic development. Lateral line was labeled with green fluorescence. Images were acquired using a 10× objective at 30□minute intervals, with 488 nm laser at 20% intensity and 200 ms exposure time. Scale bar = 100 μm. (**B**) Confocal images of a 48 hpf Tg(cldnb:lynEGFP) embryo stained for Sox2. Red, Sox2; green, lynEGFP (lateral line nerves); blue, Hoechst (nuclei). The top panel (scale bar=200 μm) and higher-magnification views of the boxed region below (30x objective; scale bar=50 μm) are presented. (**C**) Time course of actin distribution in Tg(elavl3:EGFP) embryos. Representative confocal micrographs at 3, 12, 24, and 48 hpf immunolabeled for actin. Red, actin; green, EGFP (neurons); blue, Hoechst (nuclei). For each time point, the left image is a 10x overview (scale bar=100 μm), and the right images are 30x magnifications. scale bar=50 μm.

**Fig. S9.**
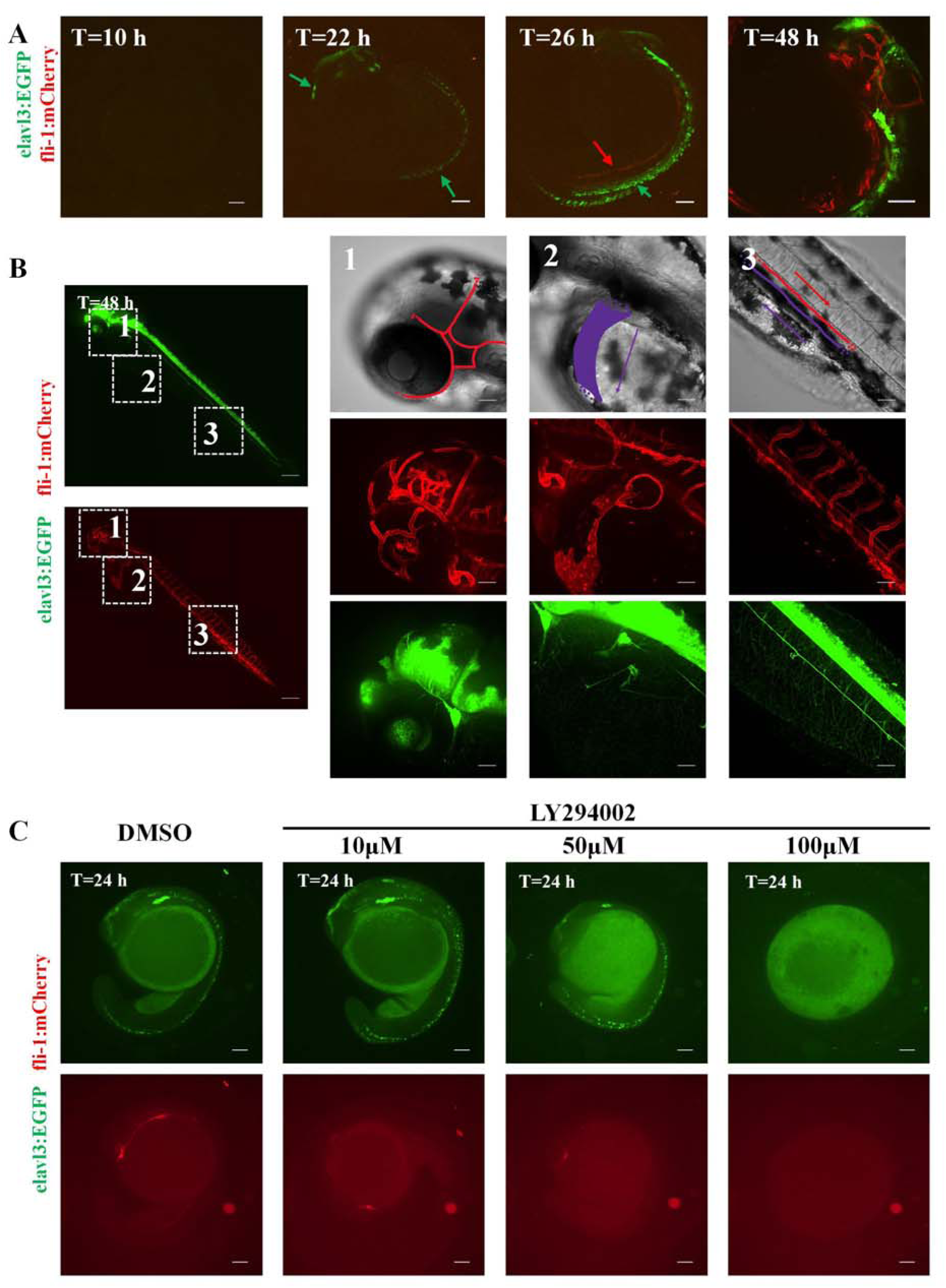
Pharmaceutical Interference on Network Formation and Coevolution of Neural Networks and Blood Vessels. (**A**) Representative confocal image of early neural and vascular development in a zebrafish embryo. Green fluorescence labels pan-neuronal structures; red fluorescence marks vascular endothelium. Green arrows indicate neuronal signals; red arrows indicate vascular signals, scale bar=100 μm. (**B**) Representative confocal images showing nervous and vascular development in a zebrafish embryo at 48h, obtained from a cross between elavl3:EGFP and fli-1:mCherry transgenic lines. The right panels display bright-field, vascular (mCherry), and neuronal (EGFP) images of the brain, yolk sac, and spinal cord regions. The leftmost confocal image was acquired with a 10x objective lens. Scale bars =200 µm. The three sets of images on the right were obtained using a 30x objective lens. Scale bars =50 µm. (**C**) Representative confocal images of zebrafish embryos following 24-hour treatment with LY294002 at concentrations of 10 μM, 50 μM, and 100 μM. The embryos are from a double transgenic Tg(elavl3:EGFP; fli-1:mCherry) line, with green fluorescence labeling neurons and red fluorescence labeling blood vessels. Scale bar = 100 μm.

**Fig. S10.**
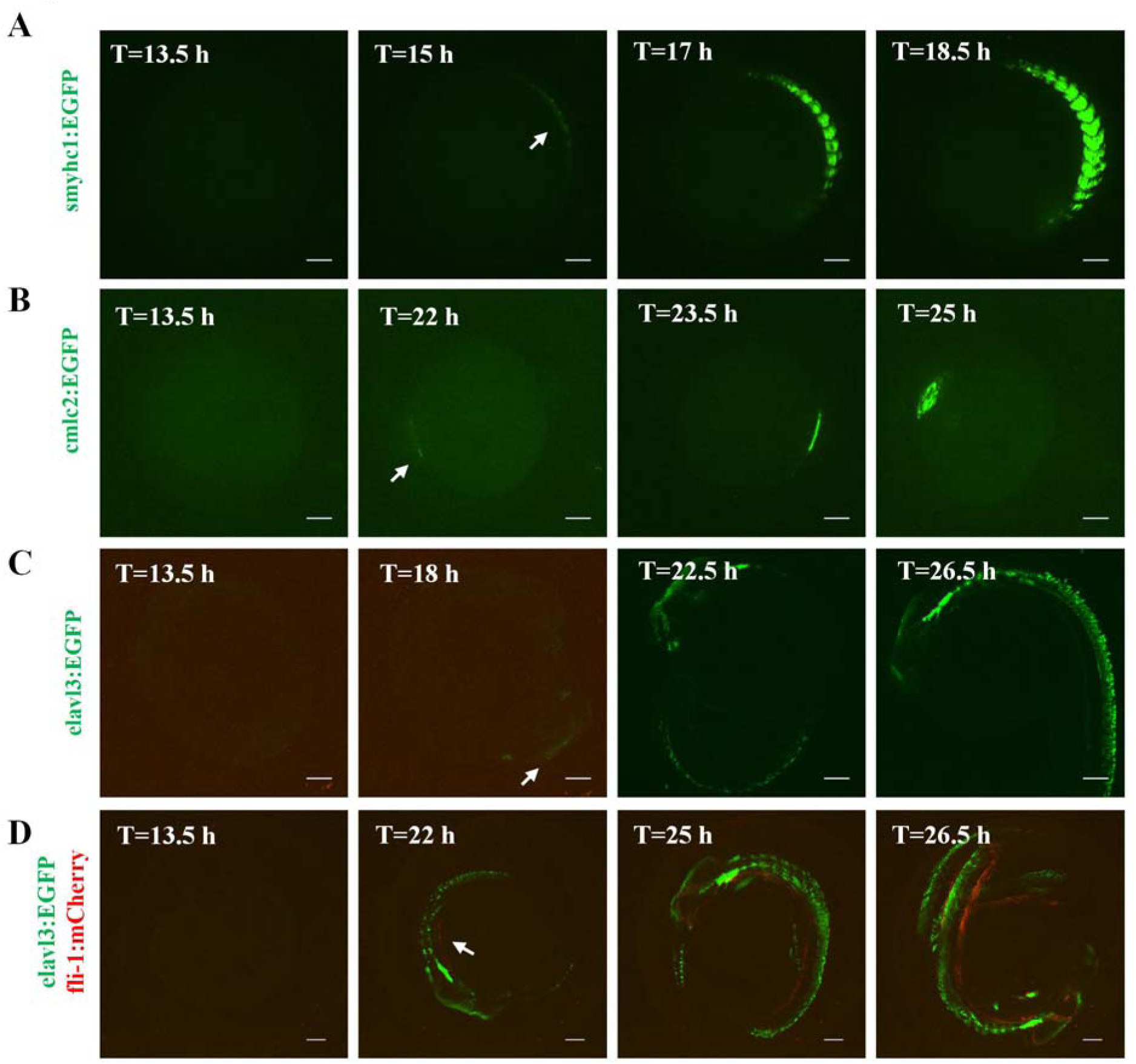
Relationship between neurogenesis and organogenesis. (**A**) Representative confocal microscopy images of live zebrafish (smyhc1:EGFP) embryos. Skeletal muscle is labeled with green fluorescent protein (GFP). The white arrow indicates the time point and location of distinct skeletal muscle signals during embryonic development. Scale bar = 100 μm. (**B**) Representative confocal microscopy images of live zebrafish (cmlc2:EGFP) embryos. Cardiac muscle is labeled with GFP. The white arrow indicates the time point and location of distinct cardiac muscle signals during embryonic development. Scale bar = 100 μm. (**C**) Representative confocal microscopy images of live zebrafish (elavl3:EGFP) embryos. Pan-neuronal structures are labeled with GFP. The white arrow indicates the time point and location of distinct neuronal signals during embryonic development. Scale bar = 100 μm. (**D**) Representative confocal microscopy images of live zebrafish (elavl3:EGFP; fli-1:mCherry) embryos. Pan-neuronal structures are labeled with GFP, and vascular endothelium is labeled with mCherry. The white arrow indicates the time point and location of distinct vascular signals during embryonic development. Scale bar = 100 μm.

**Fig. S11.**
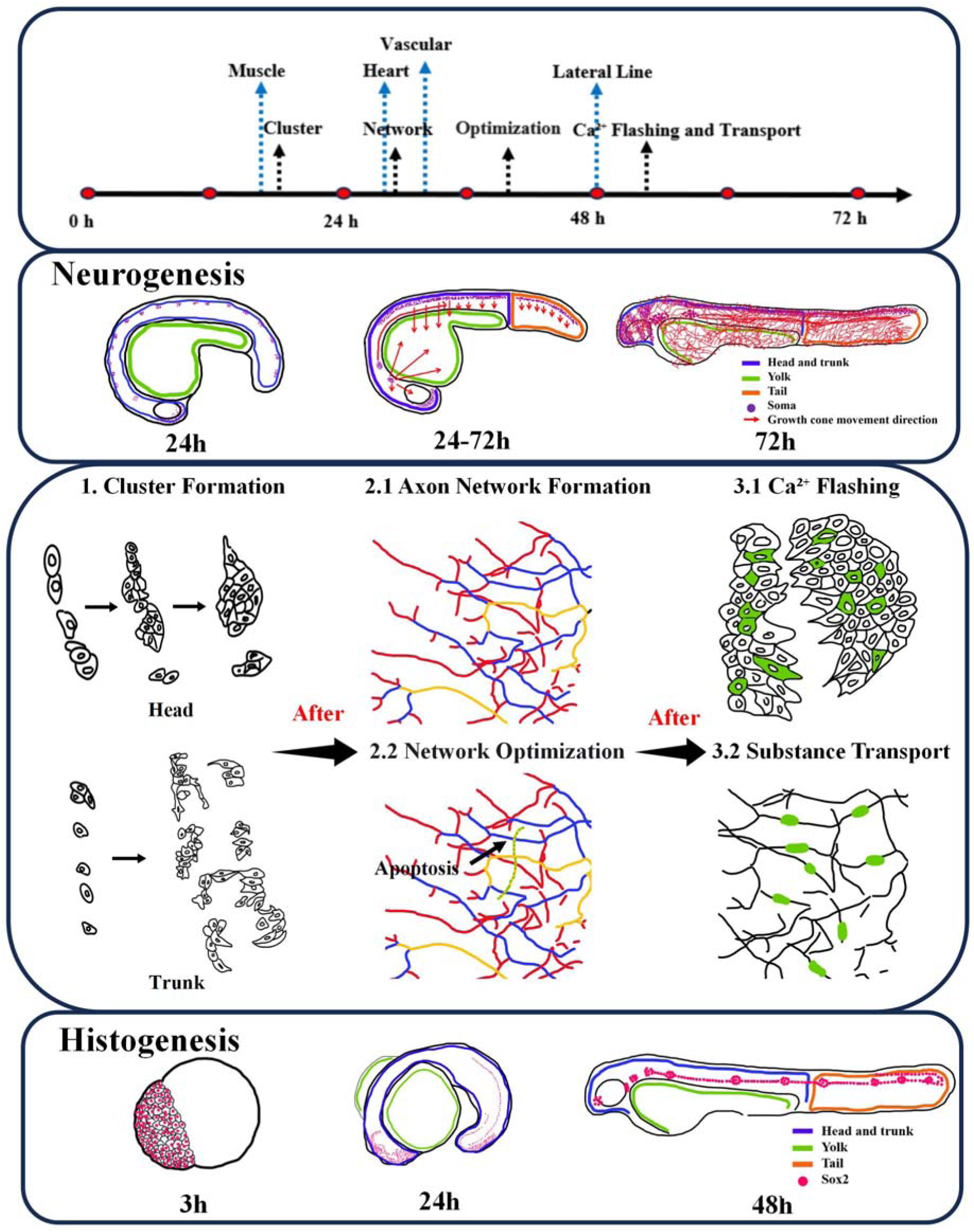
Schematic summary of the findings. In this study, we pioneered the use of live 3D imaging to reconstruct the complete timeline of neural network development and organogenesis in zebrafish. We found that neural network initiation involves the appearance of base station-like clusters in the brain and spinal cord, from which networks extend to cover the brain, spinal cord, and yolk sac. This network is subsequently refined via an apoptosis-like pruning process. Once established, the network displays frequent somatic calcium transients and active material transport in neurons.

**Table S1.** Summary of all zebrafish strains used in this study.

| ID | Name | Description | Excitation wavelength [nm] |
| --- | --- | --- | --- |
| 1 | Tg(elavl3:EGFP) | Visualize the Nervous System | 488 |
| 2 | Tg(elavl3:GCaMP6f) | Visualize of Neural Calcium Ion Signals | 488 |
| 3 | Tg(fli-1:mCherry) | Visualize Blood Vessels | 561 |
| 4 | Tg(cldnb:lynEGFP) | Visualize Lateral Line Neuromast | 488 |
| 5 | Tg(smyhc1:EGFP) | Visualize Skeletal Muscle | 488 |
| 6 | Tg(cmlc2:EGFP) | Visualize Cardiac Muscle | 488 |
| 7 | Zebrafish(AB) | Wild-Type Zebrafish | N.A |

**Table S2.** Summary of all reagents used in this work.

| ID | Name | Manufacturer | Catalog Number |
| --- | --- | --- | --- |
| 1 | LY294002 | MCE | HY-10108 |
| 2 | SiR-actin Kit | Cytoskeleton | CY-SC001 |
| 3 | Hoechst 33258 | Pumei Biological | PMK0965 |
| 4 | Anti-SOX2 | Abcam | ab97959 |
| 5 | IgG H&L (Alexa Fluor® 594) | Abcam | ab150080 |

**Table S3.** Common differentially expressed genes shared by highly expressed neural-related genes and cell death-related genes during zebrafish embryonic development (A total of 4 genes were identified. As paralogous genes, ank2a and ank2b exhibit similar functions and are therefore listed in the same column in the example table).

| ID | Name | Role in Neurodevelopment | Role in Cell Death |
| --- | --- | --- | --- |
| 1 | ank2a/b | Organizes the Axon Initial Segment (AIS) and nodes of Ranvier. | Loss of structural anchorage triggers "anoikis-like" programmed cell death. |
| 2 | ngfrb | Guides neural crest migration and axonal pathfinding. | Activates pro-apoptotic JNK signaling in the absence of neurotrophic support. |
| 3 | nradd | Modulates the affinity of neurotrophin receptors. | Directly recruits death-domain proteins to initiate the Caspase cascade. |

**Table S4.** Common differentially expressed genes shared by highly expressed neural-related genes and calcium ion-related genes during zebrafish embryonic development (A total of 20 genes were analyzed, cacna2d1a, cacna2d2b, cacna2d3, cacna2d4a, and cacna2d4b, all belonging to the cacna2d gene family and encoding voltage-gated calcium channel α□δ auxiliary subunits, are listed in the same column in the example table, ncs1a and ncs1b are paralogous genes, and caln1 and caln2 are orthologous subtypes; both are involved in the regulation of synaptic calcium signaling and are listed in the same column, notch1a, notch1b, notch2, and notch3 are all Notch signaling receptor genes in zebrafish and are placed in the same column, nell2a and nell2b are paralogous genes and are listed in the same column.).

| ID | Name | Role in Neurodevelopment | Role in Calcium |
| --- | --- | --- | --- |
| 1 | cacna2d1a/2d2b<br>/2d3/2d4a/b | Presynaptic assembly;<br>neurotransmitter release. | Enhances Ca<sup>2+</sup> current<br>amplitude and kinetics. |
| 2 | ncs1a/b, caln1/2 | Growth cone navigation; synaptic<br>plasticity. | Senses intracellular Ca<sup>2+</sup><br>fluctuations; activates<br>phosphatases. |
| 3 | notch1a/b,<br>notch2, notch3 | Lateral inhibition; maintenance of<br>neural progenitor pool. | Ca<sup>2+</sup>-dependent regulation<br>of receptor proteolysis. |
| 4 | nell2a, nell2b | Promotes axonal fasciculation and<br>hippocampal development. | Modulates ER Ca<sup>2+</sup> release<br>and steady-state levels. |
| 5 | s100b | Astrocyte marker; promotes<br>neurite extension at nanomolar<br>levels. | Extracellular Ca<sup>2+</sup>-binding<br>protein that acts as a<br>localized signaling cytokine. |
| 6 | ppef1 | Specialized in the development of<br>sensory neurons (e.g., lateral line). | A phosphatase directly<br>regulated by<br>Ca<sup>2+</sup>/Calmodulin binding. |

**Table S5.**
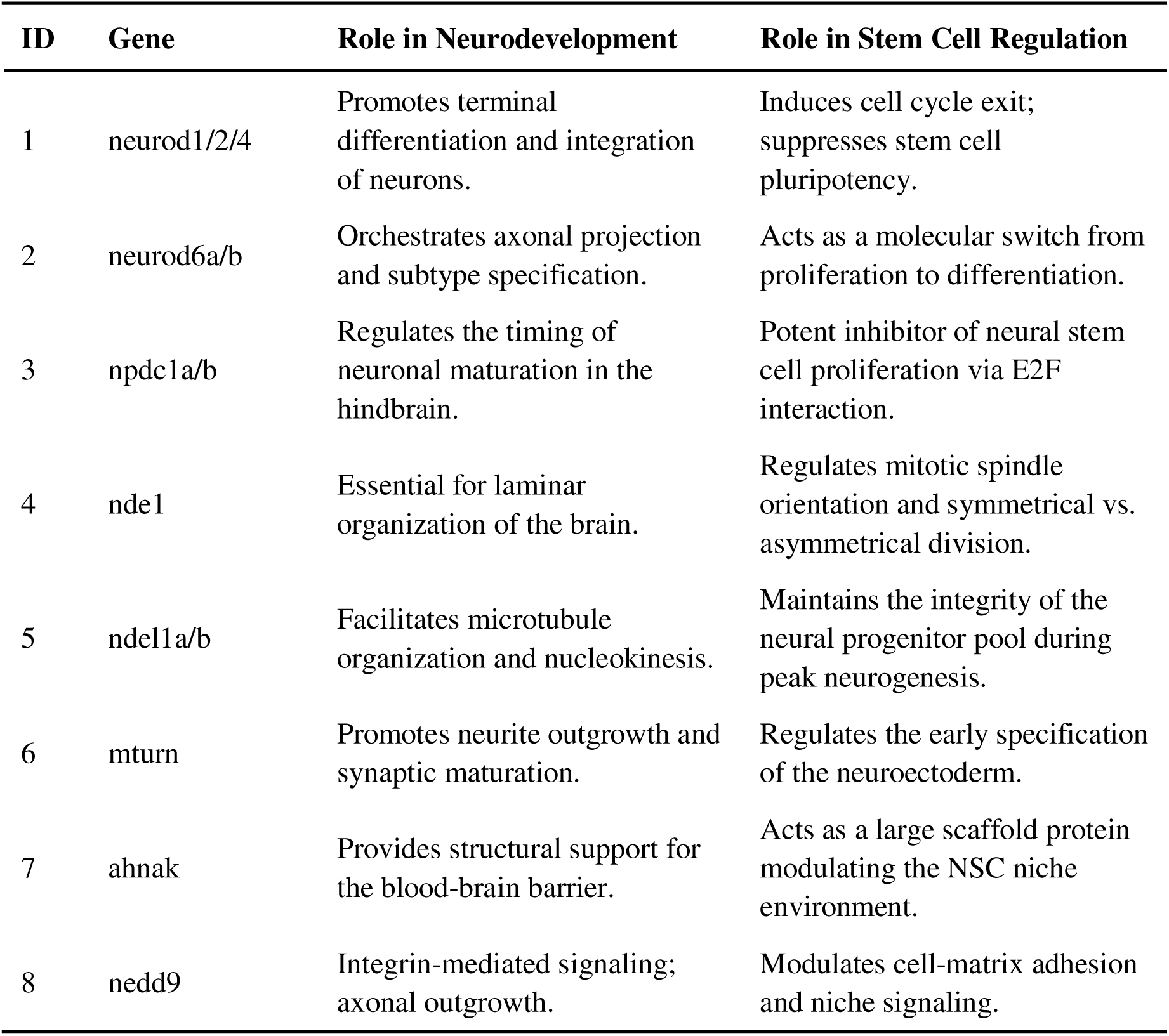
Common differentially expressed genes shared by highly expressed neural-related genes and stem cell-related genes during zebrafish embryonic development (A total of 14 genes were analyzed, neurod1/2/4 belong to the NeuroD family and collectively drive neuronal differentiation, and are thus listed in the same column in the example table, neurod6a and neurod6b, as members of the NeuroD family, regulate neurogenesis and are placed in the same column, nnpdc1a and npdc1b are both membrane-associated proteins containing the conserved NPDC1 domain, with high sequence homology, and are listed in the same column, ndel1a and ndel1b are both microtubule-binding proteins harboring the conserved NudE domain, and are grouped in the same column.).

**Table S6.** This table summarizes the gene functions of specific promoters in distinct transgenic zebrafish lines and the developmental stages of zebrafish where corresponding fluorescent signals were detected, as well as the gene functions targeted by antibodies used for immunofluorescence staining and the relevant zebrafish developmental stages with detectable fluorescent signals.

| ID | Gene | Biological Function | Signal onset time | Method |
| --- | --- | --- | --- | --- |
| 1 | sox2 | sox2 is a SOX family transcription factor and a conserved core stem cell marker in vertebrates, with dual core functions in pluripotency maintenance and neural development regulation. It mainly maintains the self-renewal and pluripotent differentiation potential of embryonic and induced pluripotent stem cells, and prevents premature stem cell differentiation; meanwhile, it regulates neural plate induction, neural progenitor proliferation and neurogenesis, acting as a key regulator for sustaining neural stem cell stemness. This gene is highly and specifically expressed solely in stem cells, neural stem cells and neural progenitors, but silenced in differentiated mature somatic cells. | <12 hpf<br>(label embryonic stem cells)<br><24 hpf<br>(neural stem cells) | Immunofluorescent staining (sox2) |
| 2 | smyhc1 | smyhc1 refers to slow myosin heavy chain 1 gene, which encodes the myosin heavy chain essential for slow muscle contraction, regulates sarcomere assembly and slow muscle function, and is specifically expressed in slow skeletal muscle. | ~ 15 hpf | Live cell imaging (smyhc1:EGFP) |
| 3 | elavl3 | elavl3 is an RNA-binding protein gene of the ELAVL family. It regulates gene expression at the post-transcriptional level, stabilizes a series of mRNAs critical for neuronal identity and function, and is exclusively expressed in neurons. | ~ 18 hpf | Live cell imaging (elavl3:EGFP) |
| 4 | fli-1 | fli1 is a key transcription factor of the ETS family that regulates vascular endothelial cell differentiation and angiogenesis during embryonic development. | ~ 22 hpf | Live cell imaging (fli-1:mCherry) |
| 5 | cldnb | cldnb refers to claudin b gene, a member of the claudin tight junction protein family. It participates in the assembly and maintenance of intercellular tight junctions, regulates cell polarity, cell migration and tissue barrier formation, and is highly and specifically expressed in lateral line primordium, neuromasts and other lateral line-related structures in zebrafish. | <24 hpf<br>~ 48 hpf<br>(label posterior lateral line) | Immunofluorescent staining and live cell imaging (cldnb:lynEGFP) |
| 6 | cmlc2 | cmlc2 refers to cardiac myosin light chain 2 gene, belonging to the myosin light chain family and serving as a hallmark gene of zebrafish cardiac muscle. It encodes the myosin light chain essential for cardiac contraction, participates in cardiac sarcomere assembly and the regulation of cardiomyocyte contraction and relaxation, maintains normal cardiac rhythm and pumping function, and is exclusively expressed in cardiomyocytes without ectopic expression in other tissues. | ~ 22 hpf | Live cell imaging (cmlc2:EGFP) |
| 8 | SiR actin | actin refers to the actin gene, encoding highly conserved cytoskeletal proteins. | <3 hpf | Immunofluorescent staining (SiR actin) |

**Movie S1.**

**Two-photon imaging and 3D reconstruction of the nervous system in a zebrafish (elavl3:EGFP) embryo.** A 30 hpf zebrafish embryo was immobilized in low-melting-point agarose and imaged with a two-photon microscope (FVMPE-RS) equipped with a 25x water-immersion objective. Z-stacks were acquired with a 5 μm step size.

**Movie S2.**

**Wide-field time-lapse confocal imaging of initial neural development in a zebrafish (elavl3:EGFP) embryo.** Images were captured using a 30x silicon oil-immersion objective with tiling. Pan-neuronal structures are labeled with EGFP (excited at 488 nm; laser power 20%; exposure time 300 ms). Imaging started at 12 hpf with a 30-minute interval over 16.5 hours. Scale bar = 200 μm.

**Movie S3.**

**Wide-field time-lapse confocal imaging of neural network formation in the brain and yolk sac of a zebrafish (elavl3:EGFP) embryo.** Images were acquired using a 30x silicon oil-immersion objective. Pan-neuronal structures are labeled with EGFP (488 nm excitation; 20% laser power; 200 ms exposure). Time-lapse imaging started at 24 hpf with a 20-minute interval for over 38 hours. Scale bar = 50 μm.

**Movie S4.**

**Movie S4. Wide-field time-lapse confocal imaging of major ventral neuron growth toward the spinal cord(left) in zebrafish (elavl3:EGFP) embryos and lateral line growth(right) in zebrafish (cldnb:lynEGFP) embryos.** Images were acquired using a 10x objective. The left video shows wide-field time-lapse confocal imaging of major ventral neuron growth toward the spinal cord in zebrafish (elavl3:EGFP) embryos. Pan-neuronal structures were labeled with EGFP (excitation: 488 nm, laser power: 20%, exposure time: 300 ms). Imaging started at 24 hpf, with 20-minute intervals for more than 22 hours. The right video shows wide-field time-lapse confocal imaging of lateral line growth in zebrafish (cldnb:lynEGFP) embryos. Lateral line structures were labeled with lynEGFP (excitation: 488 nm, laser power: 20%, exposure time: 200 ms). Imaging started at 34 hpf, with 30-minute intervals for more than 10 hours. Scale bar = 100 μm.

**Movie S5.**

**High-magnification time-lapse confocal imaging of neural network formation in a zebrafish (elavl3:EGFP) embryo.** Images were acquired using a 60x oil-immersion objective. Pan-neuronal structures are labeled with EGFP (488 nm excitation; 20% laser power; 200 ms exposure). Imaging started at 24 hpf with a 20-minute interval over 47 hours. Scale bar = 20 μm.

**Movie S6.**

**A Wide-field time-lapse confocal imaging of neural network formation in the spinal cord of a zebrafish (elavl3:EGFP) embryo.** Images were acquired using a 30x silicon oil-immersion objective. Pan-neuronal structures are labeled with EGFP (488 nm excitation; 20% laser power; 200 ms exposure). Imaging started at 24 hpf with a 10-minute interval for over 8.5 hours. Scale bar = 50 μm.

**Movie S7.**

**Wide-field time-lapse confocal imaging of neural network refinement in a zebrafish (elavl3:CaMP6f) embryo.** Images were acquired using a 30x silicon oil-immersion objective. Calcium signals in pan-neuronal structures are indicated by CaMP6f (488 nm excitation; 20% laser power; 200 ms exposure). The left video started at 33.5 hpf with a 25-minute interval; the right video started at 50 hpf with a 10-minute interval. Scale bar = 50 μm.

**Movie S8.**

**Wide-field time-lapse confocal imaging of somatic calcium transients in the brain of a zebrafish (elavl3:CaMP6f) embryo.** Images were acquired using a 30x silicon oil-immersion objective. Calcium signals in pan-neuronal structures are indicated by CaMP6f (488 nm excitation; 20% laser power; 200 ms exposure). The left video started at 24 hpf; the right video started at 52 hpf. Both were acquired with a 0.491-second interval. Scale bar = 50 μm.

**Movie S9.**

**Wide-field time-lapse confocal imaging of neural transport in zebrafish embryos.** Images were acquired using a 30x silicon oil-immersion objective (488 nm excitation; 20% laser power; 200 ms exposure). Left: Zebrafish (elavl3:CaMP6f) showing calcium signals in pan-neuronal structures. Imaging started at 50 hpf with a ∼4-second interval. Right: Zebrafish (elavl3:EGFP) with EGFP-labeled pan-neuronal structures. Imaging started at 48 hpf with a ∼10-minute interval. Scale bar = 50 μm.

**Movie S10.**

**Time-lapse light-field imaging of the angeiogenesis process in a zebrafish (elavl3:EGFP; fli-1:mCherry) embryo.** Images were acquired using a 40× water-immersion objective with volumetric imaging at an imaging depth of 110 μm. Pan-neuronal structures are labeled with EGFP, and vascular structures are labeled with mCherry. Time-lapse imaging started at 24 hpf with a 10-minute interval for over 20 hours.

**Movie S11.**

**Time-lapse confocal imaging of blood cell circulation in a zebrafish embryo.** Images were acquired using a 30x silicon oil-immersion objective. Imaging started at 24 hpf with a 1-second interval. The left panel shows blood flow in the brain, the middle panel shows circulation on the yolk sac surface, and the right panel shows spinal cord blood flow. Scale bar = 50 μm.

